# Sorting liposomes of distinct sizes by DNA-brick assisted centrifugation

**DOI:** 10.1101/2020.02.01.930321

**Authors:** Yang Yang, Zhenyong Wu, Laurie Wang, Kaifeng Zhou, Kai Xia, Qiancheng Xiong, Yong Xiong, Thomas J Melia, Erdem Karatekin, Hongzhou Gu, Chenxiang Lin

## Abstract

The “tiny bubbles of fluid” wrapped by lipid-bilayer membranes, termed vesicles, are abundant in cells and extracellular space, performing critical tasks including nutrient uptake, cargo transport, and waste confinement. Vesicles on different missions and transport routes are often distinct both in size and in chemical composition, which confers specificity to their interactions with other membranous compartments. Therefore, to accurately recapitulate the vesicles’ structure and behavior, it is important to use homogeneous liposomes (vesicles made of synthetic components) with precisely defined attributes as model membranes. Although existing methods can generate liposomes of selected sizes with reasonable homogeneity, the scalable production of uniformly-sized liposomes across a wide range of dimensions and compositions remains challenging. Here we report a streamlined, high-throughput sorting technique that uses cholesterol-modified “nanobricks” made of a few DNA oligonucleotides to differentiate hetero-sized liposomes by their buoyant densities. After DNA-brick coating, milligrams of liposomes of different origins (e.g., produced via extrusion or sonication, and reconstituted with membrane proteins) can be separated by centrifugation into six to eight homogeneous populations with mean diameters from 30 to 130 nm. In proof of concept experiments, we show that these uniform, leak-free liposomes are ideal substrates to study, with an unprecedented resolution, how membrane curvature influences the activity of peripheral (ATG3) and integral (SNARE) membrane proteins. We anticipate that our sorting technique will facilitate the quantitative understanding of membrane curvature in vesicular transport. Furthermore, adding a facile and standardized separation step to the conventional liposome preparation pipeline may benefit the formulation and prototyping of liposomal drug-carrying vehicles.

---

Classical methods for controlling liposome size rely on liposome formation conditions^1–3^ (e.g., lipid composition and solvent-to-water mixing ratio) as well as post-formation homogenization^4,5^ (e.g., extrusion and sonication) and purification^6,7^ (e.g., centrifugation and size-exclusion chromatography). The production outcome is tied to a set of empirically determined parameters that may not be independently tunable, thus limiting users’ ability to selectively vary the liposome size and composition. Microfluidic-based systems provide a way to tune liposome size and dispersity, but often require nonstandard devices built in-house^8,9^. Additionally, the capability of microfluidic-basic methods to make functional proteoliposomes is yet to be examined. Another promising approach is to guide lipid-bilayer self-assembly by DNA nanotemplates^10–12^. While effective in forming size-controlled liposomes with programmable membrane-protein stoichiometry, this approach is cost-ineffective for mass production due to the requirement of a unique DNA template for each liposome configuration and the relatively low lipid recovery. Moreover, the use of detergent limits the selection of compatible cargo molecules.

To overcome these problems, here we devise a liposome sorting strategy (**Fig. 1a**) that can be used in conjunction with an assortment of liposome manufacturing methods. Although typical lipid bilayers are lighter than aqueous solutions, liposomes that are different in size but identical in membrane and internal contents differ only slightly in buoyant density, because a liposome’s aqueous lumen constitutes the bulk of its mass. However, the surface-area-to-volume ratio (S/V) of a spherical liposome decreases rapidly with increasing size (i.e., S/V is inversely proportional to radius), affording the opportunity to amplify the buoyant density difference among liposomes by ubiquitously coating them with a dense material (similar to attaching bricks to helium balloons). In theory, smaller liposomes will gain more density than larger ones when coated by such molecular bricks (**Fig. 1b**), allowing liposome separation by isopycnic centrifugation.

**Figure 1.**
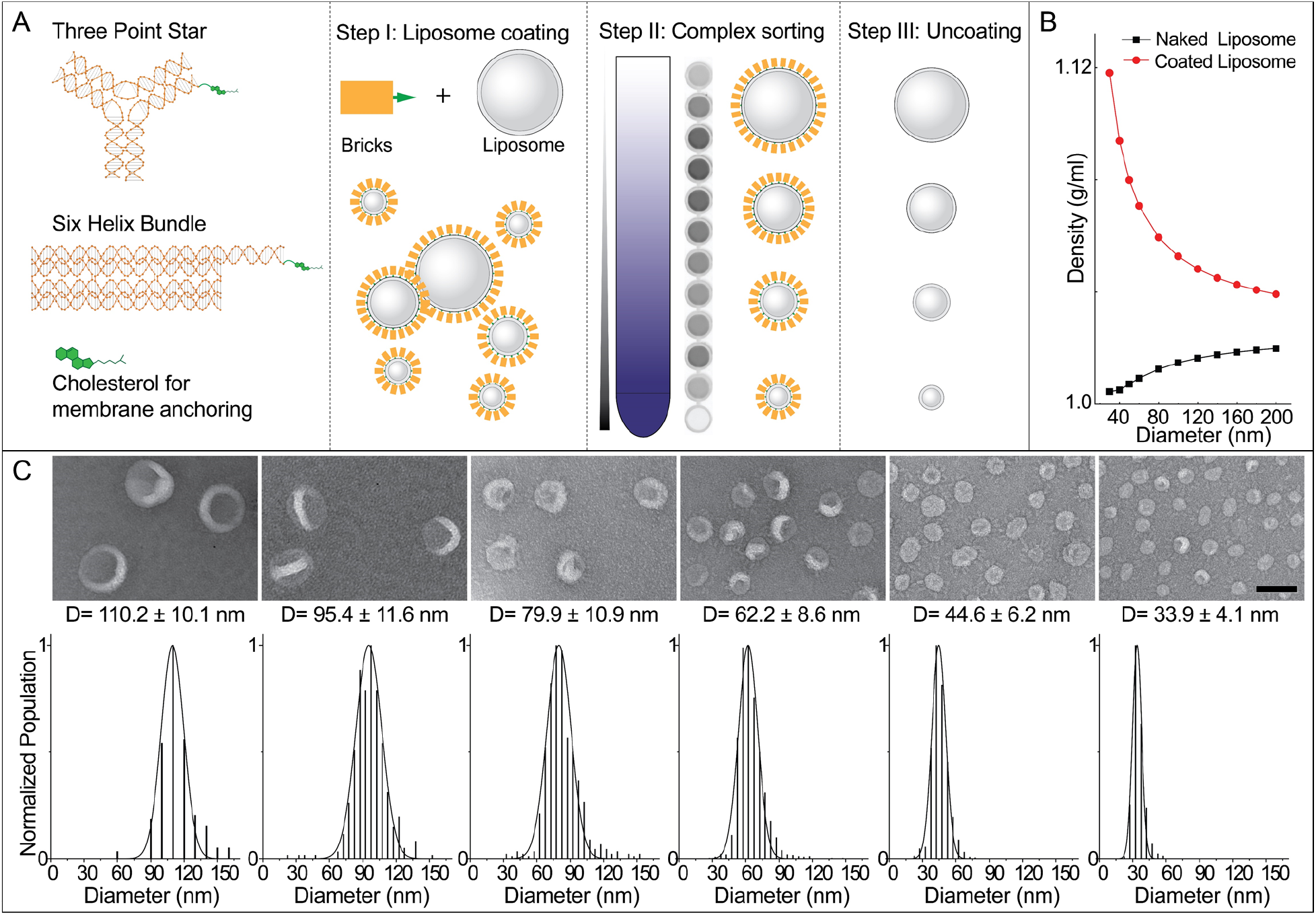
DNA-brick-assisted liposome sorting scheme and results. (A) Schematic diagrams of cholesterol-labeled DNA bricks (left) and brick-assisted liposome sorting (right) — liposome coating by DNA bricks, separation of DNA-coated liposomes by isopycnic centrifugation, and removal of DNA bricks from the sorted liposomes. A monochromic fluorescence image of 12 fractions recovered after centrifugation (Step II) shows the spread of liposomes in the density gradient. (B) A plot showing buoyant densities of naked and DNA-coated liposomes of various sizes. The theoretical values were calculated assuming the buoyant density, footprint, and molecular weight of a six-helix bundle DNA brick to be 1.7 g/cm^3^, 189 nm^2^ and 189 kD, respectively (see Supplementary Information for details), and only meant to illustrate the general trends of liposome density versus size in the presence and absence of DNA coating. (C) Liposomes sorted into distinct sizes (shown as D=mean±SD) with the help of the six-helix-bundle DNA bricks. Representative negative-stain TEM images are shown above the corresponding histograms (N=156–1690) fitted by Gaussian functions. Liposomes are made of ~59.2% 1,2-dioleoyl-sn-glycero-3-phosphocholine (DOPC), 30% 1,2-dioleoyl-sn-glycero-3-phosphoethanolamine (DOPE), 10% 1,2-dioleoyl-sn-glycero-3-phospho-L-serine (DOPS), and 0.8% 1,2-dioleoyl-sn-glycero-3-phosphoethanolamine-N-(lissamine rhodamine B sulfonyl) (rhodamine-DOPE). Scale bar: 100 nm.

We chose DNA as the coating material for its high buoyant density (~1.7 g/mL in CsCl medium)^13^, excellent solubility, programmable self-assembly behaviors^14^, and easiness to conjugate with hydrophobic molecules^15^. Previously, designer DNA nanostructures bearing hydrophobic moieties have shown promise in functionalizing and deforming liposomes^16–18^. In this work, we built two DNA structures (**Fig. 1a** and **Fig. S1**), a three-pointed star^19^ (~86 kD) and a six-helix-bundle rod^20^ (~189 kD), with a single cholesterol at the end of each DNA structure as the membrane anchor. Placing only one hydrophobic molecule per structure minimizes the brick’s footprint on the liposome surface and limits aggregation and membrane deformation. To facilitate analysis, we labeled ~10% of DNA bricks with Cy5 fluorophore. After assembling the cholesterol-modified DNA bricks by thermal annealing and purifying them by rate-zonal centrifugation (**Fig. S2**), we incubated them with a mixture of extruded and sonicated liposomes (59.2% DOPC, 30% DOPE, 10% DOPS, and 0.8% rhodamine-DOPE, see Supplementary Materials) at the brick:lipid molar ratio of 1:375. Centrifuging these DNA-coated liposomes in a gradient of isosmotic density medium (0%-22.5% iodixanol, ~5 mL per tube) at a maximum of ~300k-rcf for 4.5 hours spread the liposomes into a smeared band spanning the central two-thirds of the gradient. Analyzing the gradient fractions (~200 μL each, named F1–F24 from top to bottom) by SDS-Agarose gel electrophoresis confirmed the coexistence of DNA bricks and liposomes in the middle portion of the gradient, and revealed free DNA bricks at the very bottom, suggesting the bricks may have saturated the surface of liposomes (**Fig. S3**). Negative-stain transmission electron microscopy (TEM) study showed that F6-F18 each contained uniform-size liposomes with coefficient of variation less than 15% (**Fig. 1c** and **Fig. S4**), on par with the size homogeneity achieved through DNA-template guided lipid self-assembly. This finding was corroborated by cryo-electron microscopy (cryo-EM), which further showed 77% of liposomes as unilamellar (**Fig. S5**). The multi-lamellar liposomes were most likely generated when extruding liposomes through filters with 200-nm pores^4^ before sorting. Importantly, the recovered fractions contained liposomes with quasi-continuous mean diameters in the range of 30–130 nm (larger liposomes found in lighter fractions), allowing us to select or bin any fractions for particular liposome sizes needed in downstream applications. By and large, coating liposomes with the two types of DNA bricks yielded comparable separation resolutions, while uncoated liposomes remained inseparable after centrifugation (**Fig. 1c** and **Fig. S6**). The heavier rod-shaped brick performed better when used to sort the >100-nm liposomes and the three-pointed-star brick led to a finer separation of liposomes smaller than 40 nm. The separation resolution and recovery yield (typically >90%) were consistent from batch to batch, at different separation scales (11 μg – 1.3 mg), and across a spectrum of lipid compositions, as long as the liposome surface was not overcrowded with polyethylene glycol (**Fig. S7**-**S8**). Additionally, the dense layer of DNA bricks (clearly visible by electron microscopy in the case of six-helix bundle rods) prolonged the shelf life of sorted liposomes (up to 20 weeks at room temperature, **Fig. S9**) and was readily removable by DNase I digestion (**Fig. S10**).

The well-maintained monodispersity after long-term storage and the clear, intact boundaries observed by cryo-EM were promising signs of membrane integrity of sorted liposomes. To confirm this, we used 6-helix-bundle bricks to assist the sorting of extruded liposomes (a 1:1 mixture of liposomes passed through filters with 200-nm and 50-nm pores) loaded with fluorescein-labeled class I deoxyribozymes (I-R1a), which self-cleave in minutes upon exposure to ~1 mM Zn^2+^ at near-neutral pH (**Fig. 2a**)^21^. Similar to the plain liposomes, most deoxyribozyme-loaded liposomes with DNA-brick coatings were sorted into six homogeneous populations with mean diameters from 64 to 129 nm (**Fig. 2b** and **Fig. S11**, few smaller liposomes recovered due to their scarcity in the extruded liposomes). The narrow size distribution of each sorted fraction contrasts with the heterogeneous populations generated by filter-driven homogenization (**Fig. S7**), again highlighting the effectiveness and necessity of DNA-assisted sorting. The molar ratio between lipid and deoxyribozyme (determined by the fluorescence of rhodamine and fluorescein) was inversely proportional to liposome diameter, as expected from S/V of a sphere, indicating the unbiased cargo load in all sizes of liposome (**Fig. 2c**). Moreover, the liposomes, sorted or not, were impermeable to Zn^2+^ (2 mM) and deoxyribozyme (1 μM), showing no detectable I-R1a self-cleavage when incubated with Zn^2+^-containing solutions for over 12 hours, until we lysed liposomes with detergent (1% octyl β-D-glucopyranoside).

**Figure 2.**
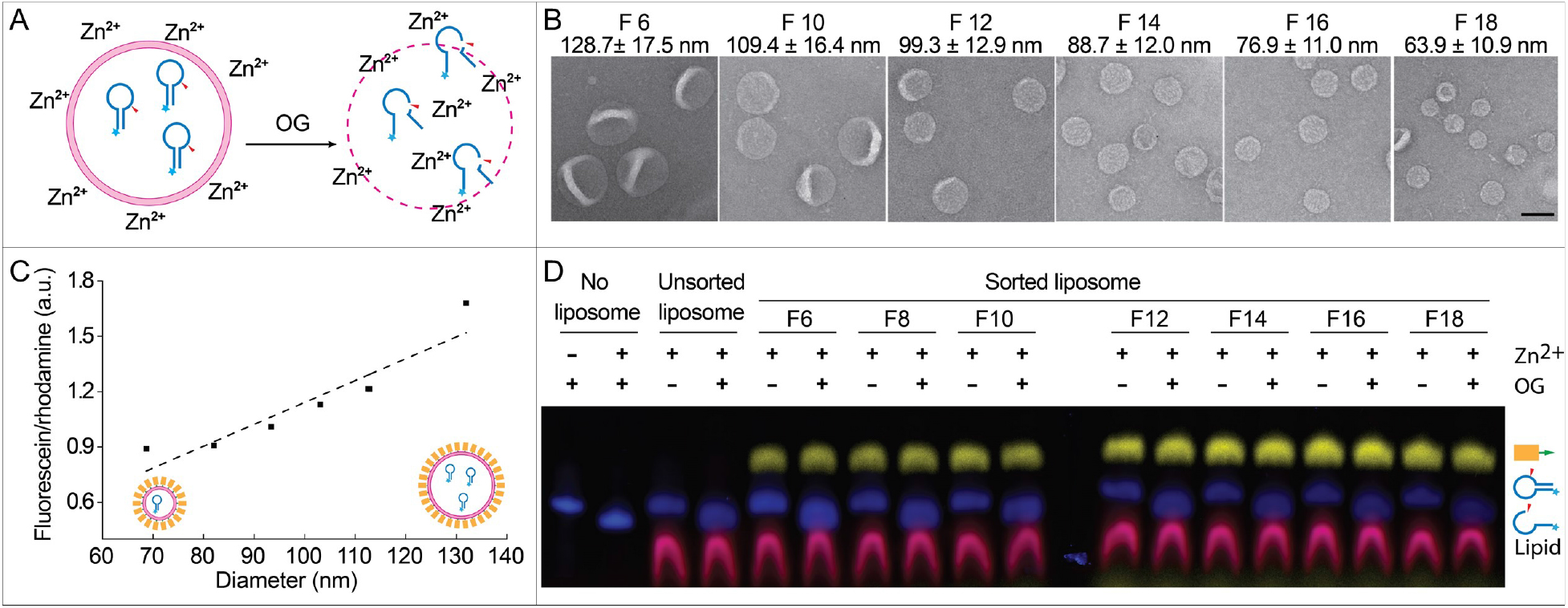
Sorting liposomes containing self-cleaving deoxyribozymes. (A) A schematic drawing of the leakage assay used to assess membrane permeability. Fluorescein-labeled deoxyribozymes undergo site-specific hydrolysis when exposed to Zn^2+^ outside of the liposomes. (B) Representative TEM images of sorted liposomes containing deoxyribozymes. Fraction numbers (e.g. F6) and liposome diameters (mean±SD, N=131–621) are noted above the corresponding images. Scale bar: 100 nm. (C) A plot showing the lipid-to-deoxyribozyme ratios in sorted liposomes fitted via linear regression (dashed line). (D) Permeability of liposomes characterized by SDS-PAGE gel electrophoresis following the deoxyribozyme-based leakage assay. Pseudo-colors: Cy5 (on DNA bricks) = yellow; fluorescein (on deoxyribozymes) = blue; rhodamine (on liposomes) = magenta. Liposomes are made of 59.2% DOPC, 30% DOPE, 10% DOPS, and 0.8% rhodamine-DOPE.

In cells, membranes are shaped into various curvatures that localize biochemical reactions and modulate membrane remodeling. Liposomes with a fine gradient of sizes provide an ideal platform to study such curvature-dependent activities *in vitro* in a systematic and precise manner. Here we applied the liposome size sorting technique to revamp two classical assays, highlighting the benefit of using uniform-size liposomes for the experimental modeling of lipid biochemistry and membrane dynamics.

We first studied the curvature-sensing capability of a conjugating enzyme that works on the membrane surface of the autophagosome. As the autophagosome grows, GABARAP-L1 (GL1) and its homologs become covalently attached to phosphatidylethanolamine lipids on the membrane surface through the serial actions of the ATG7 and ATG3 enzymes^22^. ATG3 catalyzes the final step in this cascade and its activity depends upon an amphipathic helix that senses lipid packing defects in highly curved membranes, suggesting that this protein may specifically target the rim of the cup-shaped autophagosome as a unique intracellular morphology. Previous in vitro studies revealed a curvature dependence of ATG3 activity (higher activities with 30 nm diameter liposomes than 800 nm ones)^23^, but with extruded liposome preparations and/or sonication, it was not possible to collect curvature sensing information across the biologically relevant range of 25–60 nm, where vesicles, tubules and the autophagic rim are found. Using sorted liposomes (59.2% DOPC, 30% DOPE, 10% DOPS, and 0.8% rhodamine-DOPE) of eight selected sizes (mean diameter: 30, 40, 55, 77, 90, 98, 105, and 122 nm) for ATG3-catalyzed reactions, we confirmed that the lipidation of GL1 in general favored smaller liposomes possessing higher curvature. Specifically, our data revealed a circa 5× enrichment of GL1-PE conjugates in liposomes that are 30-55 nm in diameter comparing to larger liposomes, with the lipidation peaking on liposomes with ~40-nm diameter (**Fig. 3** and **Fig. S12**–**S13**). This curvature range is reminiscent of the typical autophagosome rim (20–50 nm lamellar spacing)^22^, the inferred hotspot of ATG3-dependent lipidation *in vivo*. As ATG3 is a peripheral protein, it must gain access to the membrane surface, and thus a potential concern of using sorted liposomes is that the DNA bricks might directly impede lipidation. Though the DNA bricks are essentially inert with respect to protein activity, we assured that the membrane surface was not obscured by treating the sorted liposomes with nuclease before the lipidation assay. Overall, homogeneous liposomes improved the precision of the *in vitro* lipidation assay, enabling a quantitative measurement of the curvature-dependent ATG3/ATG7 ligation cascade.

**Figure 3.**
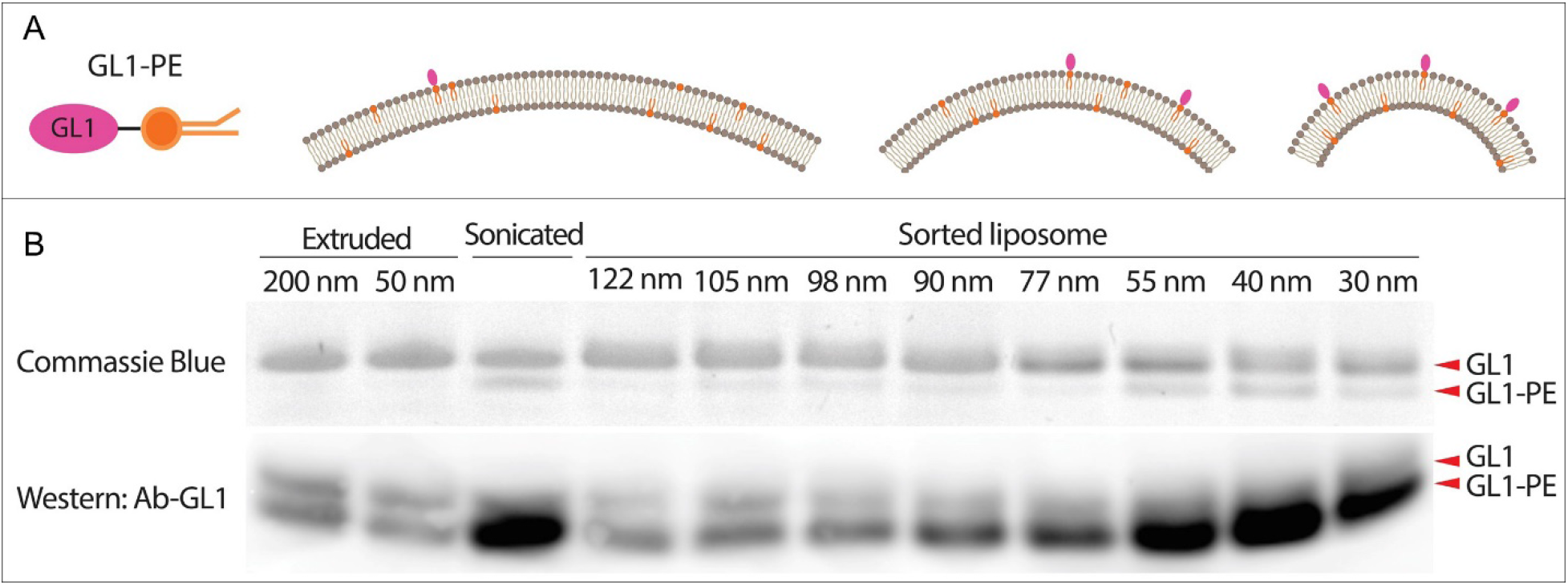
Atg3-catalyzed GL1 lipidation reaction studied using uniform-size liposomes. (A) Schematic illustrations of GL1-DOPE conjugate (left) and the expected lipidation outcomes on liposomes with differential membrane curvatures (right). (B) GL1-lipidation efficiencies on extruded, sonicated and sorted liposomes (59.2% DOPC, 30% DOPE, 10% DOPS, and 0.8% rhodamine-DOPE) characterized by gel electrophoresis (top row, stained by Coomassie Blue) and immunoblot against GL1 with an antibody that preferentially recognizes the GL1-PE conformation (bottom row). The numbers (in nm) above lanes represent the nominal pore size of the filters (extruded liposomes) or measured mean diameters (sorted liposomes).

We next turned our attention to how DNA-brick mediated sorting might work with transmembrane proteins. Soluble NSF attachment protein receptors (SNAREs) are a family of proteins that fuel membrane fusion in many intracellular trafficking routes, including the vesicular release of neurotransmitters and hormones^24–26^. Two types of SNAREs, v-SNAREs on the vesicle and t-SNAREs on the target membrane, assemble into a four-helical complex to force the membranes into proximity and eventually drive fusion. Previous experimental^27,28^ and theoretical^29^ work suggests that membrane curvature may be a critical factor in determining the kinetics of fusion and the number of SNARE complexes required. However, past experiments measured the fusion rates of proteoliposomes with only one or two sizes, due to constraints in preparation of protein-reconstituted liposomes^27,28,30^. In addition, the preparation methods often produce liposomes with broad size distributions^31^. These limitations prevented systematic studies of the curvature dependence of fusion rates. Thus, it is highly desirable to develop methods that can produce proteoliposomes with sharp size distributions.

In previous work, we addressed this issue by building DNA-ring templated liposomes displaying a predetermined number of SNARE proteins^32^. Despite the uniform and controllable proteoliposome size, an exhaustive examination of the impact of membrane curvature on fusion rate was impractical, because the obligated redesign of DNA templates for each liposome size and the small preparation scale (typically less than a few micrograms) limited the throughput of our fusion assay. To address this challenge, here we applied DNA-brick assisted size-sorting to produce proteoliposomes with well-defined sizes. We reconstituted the neuronal/exocytotic v-SNARE VAMP2 into liposomes (lipid:VAMP2 ≈ 200:1) containing FRET-dye-labeled lipids (NBD- and rhodamine-DOPE) and performed DNA-brick assisted sorting on 440 μg of such proteoliposomes. The pre-existence of proteins in vesicle membranes did not compromise the separation effectiveness, as confirmed by negative-stain TEM (**Fig S14**). After enzymatic removal of DNA bricks (unnecessary in hindsight as the DNA bricks did not affect fusion, see **Fig. S15**), we mixed VAMP-embedded liposomes of eight different diameters (37-104 nm) with unlabeled (and unsorted) liposomes carrying cognate t-SNAREs in separate test tubes; the mixtures (lipid concentration = 3 mM) were kept at 4°C for 2hrs, a temperature that allows vesicle docking but no fusion (**Fig. S16**). Finally, we warmed the pre-docked liposomes to 37°C and monitored NBD fluorescence for 2 hours using a fluorescence microplate reader. Merging of liposome membranes increases the distance between NBD dyes and their rhodamine quenchers, providing a read-out of lipid mixing kinetics (**Fig. 4a**). Consistent with previous findings^24,27,30^, we showed that the membrane fusion is SNARE-dependent. However, unlike the conventional assays, our setup discerned the lipid mixing kinetics as a function of vesicle size (**Fig. 4b**). When mean v-SNARE-bearing liposome diameters were within 47-104 nm, smaller liposomes fused more rapidly, with the most and least fusogenic vesicles showing ~3-fold difference in the final NBD fluorescence. Interestingly, further decreasing liposome diameter to an average of 37 nm slowed fusion moderately. Assays with halved VAMP2 density on liposomes yielded a similar trend (**Fig. 4c** and **4d**). We note that when lipid:VAMP2 ratios were held constant, smaller liposomes tended to display fewer v-SNAREs, which may explain the slower fusion of the 37-nm liposomes comparing to the 47-nm ones. That is, there seems to be an optimal combination of SNARE copies per liposome and membrane curvature — an effect that would not have been captured without the precise control of liposome sizes.

**Figure 4.**
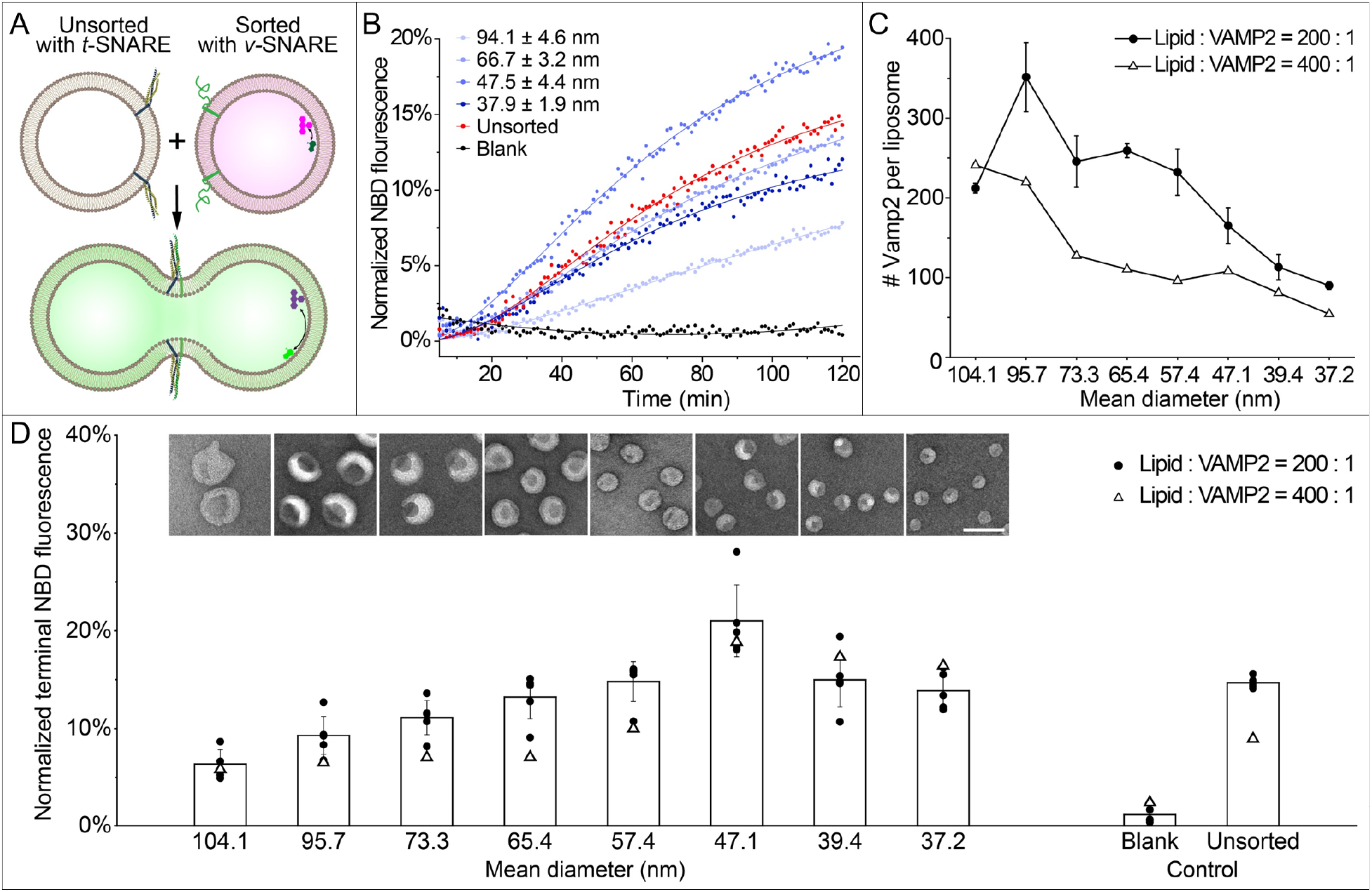
SNARE-mediated membrane fusion studied using uniform-size liposomes. (A) A schematic illustration of the lipid-mixing assay used to monitor membrane fusion. Initially quenched NBD dyes (green) fluoresce following membrane fusion due to a decrease in FRET with rhodamine dyes (magenta). SNARE proteins are shown as blue, yellow (t-SNAREs) and green (VAMP2, v-SNARE) ribbons on the membranes. (B) Representative fluorescence traces showing the kinetics of fusion between unsorted liposomes bearing t-SNAREs and unsorted (red) or sorted (different shades of blue, diameters marked as mean±SD, N>208) liposomes bearing v-SNAREs. Protein-free liposomes are mixed with v-SNARE bearing liposomes as a negative control (black). Liposomes with v-SNAREs are reconstituted with 82% POPC, 12% DOPS, 1.5% Rhodamine-DOPE, 1.5% 1,2-dioleoyl-sn-glycero-3-phosphoethanolamine-N-(7-nitro-2-1,3-benzoxadiazol-4-yl) (NBD-DOPE), and a lipid:protein molar ratio of 200:1 or 400:1. Liposomes with t-SNAREs are reconstituted with 58% 1-palmitoyl-2-oleoyl-glycero-3-phosphocholine (POPC), 25% DOPS, 15% 1-palmitoyl-2-oleoyl-sn-glycero-3-phosphoethanolamine (POPE), 2% phosphatidylinositol 4,5-bisphosphate and a lipid:protein molar ratio of 400:1. (C) v-SNARE copy numbers per liposome measured from sorted liposomes reconstituted with lipid:VAMP2 molar ratios of 200:1 and 400:1. (D) Lipid mixing after 2 hours of fusion reactions (measured by NBD fluorescence, as shown in (B)) plotted against the average diameters of sorted v-SNARE-bearing liposomes (representative TEM images are shown). Means and SDs are based on the dataset of liposomes reconstituted with lipid:VAMP2 = 200:1. Scale bar: 100 nm.

In neurons, synaptic vesicle sizes are highly homogeneous and regulated^33,34^. Here we only studied the minimal fusion machinery (SNAREs) to prove the concept. However, the platform can in principle be adapted to model more physiological conditions, where additional proteins (e.g., Synaptogamin-1 or Munc18) affect the fate of vesicles.

Self-assembled DNA nanostructures have been interfaced with lipid bilayers in a number of unconventional ways towards the goal of programmable membrane engineering^16–18^. In the past, this took one of two forms. The first approach is to scaffold liposome formation with DNA templates, which excels at precision but any pre-existing membrane needs to be micellized before reassembly^10–12^. The second strategy is to reshape the membrane landscape of liposomes with DNA devices that oligomerize or reconfigure on command, which may preserve certain pre-existing membrane features (e.g. lipid composition, internal content) but the end products tend to be less homogeneous^.35–37^. By bridging this gap, the DNA-brick assisted liposome sorting method further advances the membrane engineering capability of DNA nanotechnology. Specifically, the method separates liposomes from virtually any source into a range of narrowly distributed sizes with minimal impact on the original membrane properties. Further, two DNA structures composed of a handful of oligonucleotides fulfilled various sorting tasks. The simplicity and robustness of the technique make it readily adaptable by any biochemical laboratory with access to research-grade ultracentrifuges (**Fig. S17**). Future method development will benefit from the programmability of DNA nanostructures. For example, coating liposomes with more massive DNA bricks could facilitate the separation of larger liposomes; changing cholesterol anchors to protein-specific ligands could enable the sorting of natural vesicles by their surface markers. In addition to the utilities in basic research, we envision the method (in its current or adapted forms) finding applications in biotechnology, such as in aiding the development of drug-delivering liposomes as well as isolating disease-specific extracellular vesicles.

## Acknowledgement

This work is supported by a National Institutes of Health (NIH) Director’s New Innovator Award (GM114830), an NIH grant (GM132114), and a Yale University faculty startup fund to C.L., NIH grants to T.M. (GM100930 and GM109466) and to E.K. (NS113236), and National Natural Science Foundation of China grants to H.G. (21673050, 91859104, and 81861138004)

## Author contributions

Y.Y. initiated the project, designed and performed most of the experiments, analyzed the data, and prepared the manuscript. Z.W. performed membrane fusion study and analyzed the data. L.W. performed lipidation study. K.Z. performed cryo-EM study. K.X. replicated the sorting method. Q.X. performed negative stain TEM study. Y.X. supervised the cryo-EM study and interpreted the data. T.J.M. designed and supervised the lipidation study and interpreted the data. E.K. supervised the membrane fusion study and interpreted the data. H.G. designed the liposome leakage assay, supervised replication of the sorting method, and interpreted the data. C.L. initiated the project, designed and supervised the study, interpreted the data, and prepared the manuscript. All authors reviewed and approved the manuscript.

