## Supplementary Information for "Sorting liposomes of distinct sizes by DNA-brick assisted centrifugation"

### DNA and Lipid Materials.

DNA oligonucleotides (oligos) were synthesized by Integrated DNA Technologies. Chemically modified oligos were purified via HPLC by manufacturer, while unmodified oligos were purified via PAGE in house (see **Table S1** for oligo sequences). DNA brick designs are shown in **Figure S1**, along with the PAGE analyses of the assembly products.

**Table S1.** DNA strand sequences

| **3 Point Star Brick (3PS bricks)** | |
|---|---|
| Name | Sequence |
| C | AGGCATATTGAATCGTTTACAGGATTAGTAATTAACAGCTTTAATATCATCGCCC ATCGTAGGTTTCTTGCC |
| C-Cy5 | /5Cy5/AGGCATATTGAATCGTTTACAGGATTAGTAATTAACAGCTTTAATATCATC GCCCATCGTAGGTTTCTTGCC |
| S-a | GACGACAGAGGTTGCTAGGCG |
| S-b | TTACCGTGTGTGTTAAGGTGG |
| S-c | ACCGAGCCTCCGTCAACATCG |
| E-a | CCACCTTAACACGCGATGATATTGCTGTTAATTAGGCTCGGT |
| E-b | CGATGTTGACGGACTAATCCTGTCGATTCAATATCTGTCGTC |
| E-0 | CGCCTAGCAACCTGCCTGGCAAGCCTACGATGGACACGGTAA |
| E-Chol | CGCCTAGCAACCTGCCTGGCAAGCCTACGATGGACACGGTAA/3CholTEG/ |
| **6 Helix Bundle Brick (6HB bricks)** | |
| 6hb-M0 | TTTAGTGCTACACTGTGCGTATGCGAAAACTTGCGATATGCTCCATTT |
| 6hb-M1 | TTTAGTCGAGTGAACTGTAACGTACAGGTAGATAGACTCTGTATCTTT |
| 6hb-M2 | AAATTATCTACCACAACTCACCGCCTAGCAACCTGCCTGGCAAGCCTACGATG GACACGGTAA |
| 6hb-M3 | TTTATTCGAGCATGTCAGTGGATCAATCGTGTTAGACATGACGTATTT |
| 6hb-M4 | TTTGTGGACTATATATACGTGGAACCATGAATTGGCTGAGTTTGGTTT |
| 6hb-M5 | TTTTGGTTTACTCACTATTGTCACCTTATACCACAATCAGATCCGTTT |
| 6hb-S0 | CACAGTGGATTGTGTATATATAGTCCACTACGTCACTAGGCG |
| 6hb-S1 | CAGTTCAGTCCATCTGACATGCTCGAATCCAAACTTAAACCA |
| 6hb-S2 | TTACCGTCTCGACTTGGAGCATATCGCATAGTGAGCAGCCAA |
| 6hb-S3 | CACGATTTTCCACGGTATAAGGTGACAAAGTTTTCTACGTTA |
| 6hb-S3-Cy5 | /5Cy5/CACGATTTTCCACGGTATAAGGTGACAAAGTTTTCTACGTTA |
| 6hb-S4 | TTCATGGGATCCACGTAGGCTTGCCAGGCTACCTGGCATACG |
| 6hb-S5 | CGGATCTTAGCACTGATACAGAGTCTATCAGGTTGTGTCTAA |
| M2'-Chol | GTGAGTTGTGGTAGATAATTT/3CholTEG/ |
| **Deoxyribozyme** | |
| I-R1a-FAM | /56FAM/CATGTACAGCCATAGTTGAGCATTAAGTTGAAGTGGCTGTACATG |

All lipids were purchased from Avanti Polar Lipids. For general sorting experiments, leakage assay, and lipidation assay, liposomes were prepared in Buffer X. For the vesicle fusion study, reconstituted proteoliposomes were enriched in Buffer Y before sorting (Method **7**). To avoid osmolality shock, DNA bricks were prepared in the same buffer (X or Y) as the liposomes (see Methods **1** and **2**). Lipid and buffer compositions are listed in **Table S2**.

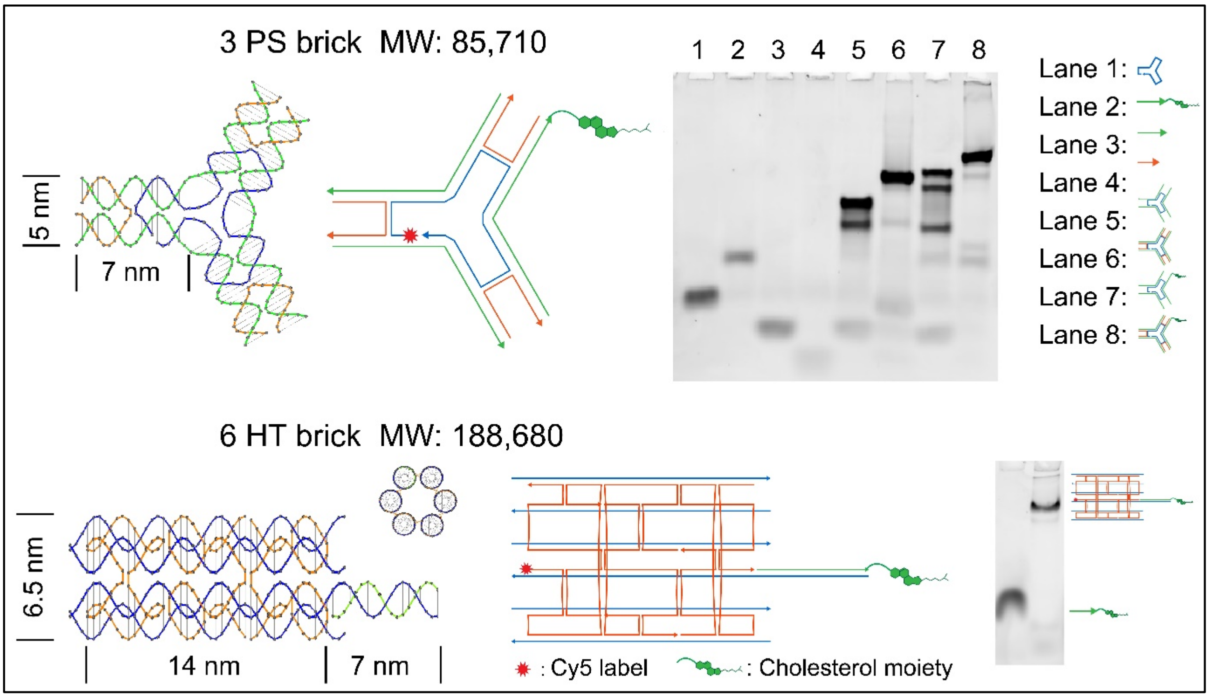

**Figure S1.** (a) 3 Point Star (3PS) and (b) 6 Helix Bundle (6HB) DNA bricks. Design diagrams are shown to the left of native PAGE results (6% gel run at 15 V/cm for 70 min).

**Table S2.** Lipid compositions and buffer ingredients. Numeric values refer to molar percentages and ratios. Composition B is used in this work unless noted otherwise.

| Abbreviation | Full name of lipids |
|---|---|
| DOPC | 1,2-dioleoyl-sn-glycero-3-phosphocholine |
| DOPE | 1,2-dioleoyl-sn-glycero-3-phosphoethanolamine |
| DOPS | 1,2-dioleoyl-sn-glycero-3-phospho-L-serine |
| DOTAP | 1,2-dioleoyl-3-trimethylammonium-propane |
| PEG-2k-DOPE | 1,2-dioleoyl-sn-glycero-3-phosphoethanolamine-N-[methoxy(polyethylene glycol)-2000] |
| rhodamine-DOPE | 1,2-dioleoyl-sn-glycero-3-phosphoethanolamine-N-(lissamine rhodamine B sulfonyl) |
| NBD-PE | 1,2-dioleoyl-sn-glycero-3-phosphoethanolamine-N-(7-nitro-2-1,3-benzoxadiazol-4-yl) |

| | DOPC | DOPE | DOPS | DOTAP | PEG-2k-DOPE | rhodamine -DOPE |
|---|---|---|---|---|---|---|
| Composition A | 99.2% | 0% | 0% | 0% | 0% | 0.8% |
| Composition B | 59.2% | 30% | 10% | 0% | 0% | 0.8% |
| Composition C | 59.2% | 30% | 0% | 10% | 0% | 0.8% |
| Composition D | 94.2% | 0% | 0% | 0% | 5% | 0.8% |

| | POPC | DOPS | rhodamine-DOPE | NBD-PE | v-SNARE:lipid |
|---|---|---|---|---|---|
| v-SNARE liposome | 82% | 15% | 1.5% | 1.5% | 1:200 or 1:400 |
| | POPC | DOPS | POPE | PIP2 | t-SNARE:lipid |
| t-SNARE liposome | 58% | 25% | 15% | 2% | 1:400 |

| | HEPES | KCl | $MgCl_2$ | pH |
|---|---|---|---|---|
| Buffer X | 25 mM | 400 mM | 10 mM | 7.0 |
| Buffer Y | 25 mM | 140 mM | 0 mM | 7.0 |

### Methods.

#### 1. Liposome preparation

*1a.* Solvent evaporation and lipid rehydration

To prepare 1 mL of liposomes containing 3 μmol total lipids (final $C_{lipid}$ = 3 mM) of a specific composition (**Table S2**), appropriate volumes of lipid stocks (dissolved in chloroform) were mixed in a round-bottom glass tube. The mixture was blown-dried under $N_2$ for at least 1/2 hour. The resulting lipid film at the tube bottom was further dried overnight in a desiccator under vacuum. To rehydrate the lipids, 1 mL of Buffer X (**Table S2**) was added into the tube and agitated for 1/2 hour. To prepare for the leakage assay, 1 mL of 1 μM FAM-modified I-R1a deoxyribozyme (dissolved in Buffer X) was used instead for rehydration. The glass tubes were wrapped with aluminum foil to reduce photobleaching of fluorescent labels.

*1b.* Sequential extrusion (to produce liposome with nominal diameters of 50–200 nm)

The rehydrated lipid suspension was transferred into a 1.5 mL centrifuge tube and thermo-cycled between a liquid-nitrogen bath and a 37°C-water bath for 5–10 times. The frozen-thawed suspension was then sequentially extruded through polycarbonate filters of nominal pore sizes of 400 nm, 200 nm and 50 nm, each time using a Mini Extruder (Avanti Polar Lipids) following manufacture's recommendation. The extruded liposomes after passing through 200-nm and 50-nm filters (typically 300 μL each) were stored at 4°C; the remaining 400 μL of liposomes was sonicated as described below.

*1c.* Sonication (to produce liposome with nominal diameters <50 nm)

Extruded liposomes (~400 μL) were sonicated using a Qsonica Q125 dip-probe sonicator for 1 min (10 cycles of 1-s on, 1-s off) while sitting on an ice-water bath.

#### 2. DNA brick preparation

*2a.* Assembly

PAGE or HPLC purified oligos were dissolved in deionized, Milli-Q water (Millipore) with concentrations normalized to 120 μM each. To assemble the 3PS and 6HB DNA bricks, various amounts of cholesterol-modified oligos (1–2.5 μM) and a stoichiometric amount of unmodified oligos (1 μM each) were mixed in Buffer X and underwent thermal annealing from 95 to 4°C (held at 95, 65, 50, 42, 37, 22, and 4°C for 5 min each). The assembly products were electrophoresed in a non-denaturing 6% polyacrylamide gel under 15V/cm for 70 min in 1×TAE, 10 mM $MgCl_2$. The optimal molar ratio between modified and unlabeled oligos, which gave rise to a sharp, distinct band after Sybr Gold staining, was chosen for DNA brick assembly for the rest of this study. Optionally, 10% of an unmodified oligo (C in 3PS and 6hb-S3 in 6HB, **Table S1**) was replaced with a Cy5-labeled oligo for staining-free visualization of DNA bricks on gels.

*2b.* Purification and characterization

Large scale (400 μL of 5 μM) DNA-brick assemblies were placed on top of a 5%–20% glycerol gradient in a 5-mL ultracentrifugation tube (Beckman Coulter, Cat# 344057). The sample-loaded density medium was spun at 55,000 rpm and room temperature (RT) for 4.5 hr in an SW55-Ti rotor (Beckman Coulter) before fractionated into 200-μL fractions. Five microliters of each fraction were electrophoresed in a 3.5% agarose gel containing 0.05% ethidium bromide under 10 V/cm for 1.5 hr in 0.5×TBE, 10 mM $MgCl_2$ (see an example in **Figure S2**). Fractions containing well-formed bricks (e.g., fractions 8–10 in **Figure S2**) were combined and concentrated to 50–100 μL by centrifugation (10 min at 10,000 rcf) on Amicon filtration units (Millipore) with 10 kD nominal molecular weight limit (NMWL). The concentrated sample was diluted in Buffer X or Y to 500 μL and concentrated again for a total of four times. The DNA brick concentration was determined by $OD_{260}$ measurement of a NanoDrop spectrometer

(Thermo Fisher Scientific). The purified bricks were diluted to 5 μM in Buffer X or Y and stored at -20°C.

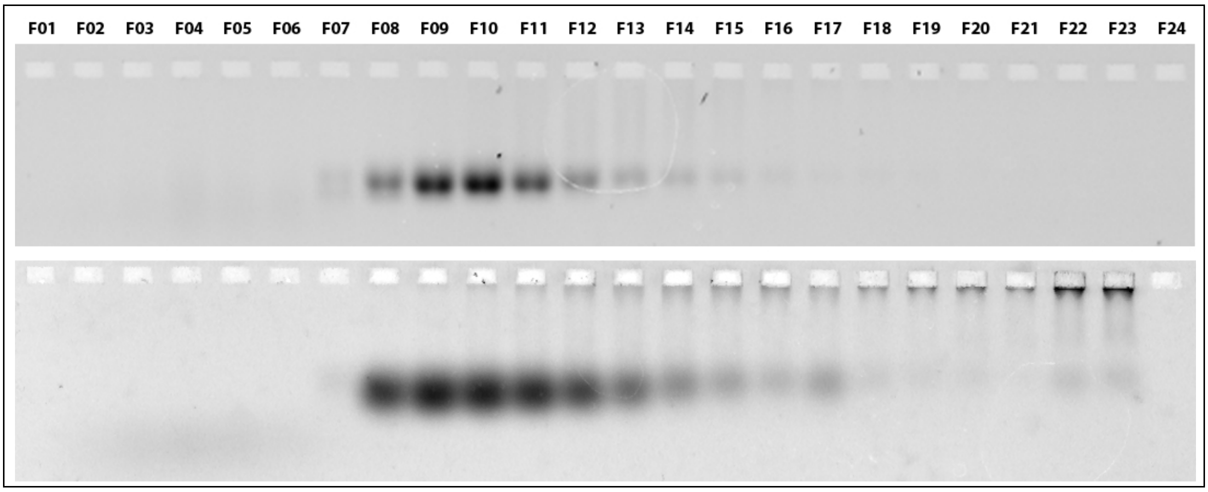

**Figure S2.** Representative images of agarose gels showing DNA bricks (top: 3PS; bottom: 6HB) after rate-zonal centrifugation. Fractions 1–24 (F01, F02 … F24) were collected from the top to the bottom of the glycerol gradient.

### 3. DNA-brick assisted liposome sorting

#### *3a.* Liposome coating

For small scale sorting, 40 μL of purified 3PS or 6HB brick (cholesterol-labeled, 1 μM) and 5 μL of liposome (3 mM lipid) were mixed in a 200 μL tube and incubated at room temperature for 1–2 hr under continuous agitation. The brick:lipid ratio of 1:375 is empirically determined to be sufficient for subsequent liposome sorting (see *3d*). In the case of suboptimal sorting, a higher concentration of DNA brick may be used for liposome coating. When sorting larger quantities of liposomes, the amount of DNA brick and liposome was increased proportionally; the DNA brick concentration may be adjusted as appropriate. **Table 3** provides some guidelines. For example, our largest scale preparation started with >1 mg liposome (1.8 μmole total lipid), which was split into six 5-mL ultracentrifuge tubes after DNA-coating for isopycnic centrifugation (see Method *3b*).

**Table S3.** The amount of reagents used for different scale of sorting experiments.

| Scale | Brick amount | Lipid amount | Total volume | Volume loaded to iodixanol gradient |
|---|---|---|---|---|
| 1× | 40 pmol | 15 nmol | 45 μL | 45 μL + 45 μL 45% iodixanol |
| 10× | 400 pmol | 150 nmol | 350 μL | 350 μL + 350 μL 45% iodixanol |
| 20× | 800 pmol | 300 nmol | 350 μL | 350 μL + 350 μL 45% iodixanol |

#### *3b.* Liposome sorting by isopycnic centrifugation

Iodixanol density gradient was prepared from stock solutions of 45%, 18%, 15%, 12%, 9%, 6%, 3% and 0% (v/v) iodixanol (Stemcell Technologies) in Buffer X.

DNA-coated liposomes were mixed with an equal volume of 45% iodixanol, forming a 22.5% iodixanol solution at the bottom of an ultracentrifugation tube. For the small scale separation (1× in **Table S3**), 80 μL of such a solution was pipetted to an 800-μL tube (Beckman Coulter Cat# 344090). Seven additional iodixanol layers (18% to 0%, 80 μL each) were carefully placed on top of one another to form a quasi-linear gradient. The tube, loaded with the liposome sample in the iodixanol gradient, was spun in an SW55-Ti rotor at 48,000 rpm and RT for 4.5 hr. For large scale preparations (e.g., 10× and 20× in **Table S3**), linear 0–18% iodixanol gradients (4.2 mL each) were formed in 5-mL tubes (Beckman Coulter, Cat# 344057) using a Gradient Master (BioComp Instruments). Seven-hundred microliters of DNA-coated liposomes in 22.5% iodixanol were carefully layered at the bottom of the gradient using a syringe and a needle. The tubes were spun at 50,000 rpm and RT for 4.5 hours. Proteoliposomes (see **Figure S14**), were sorted in the same way, except using gradients made in Buffer Y.

*3c.* Post-centrifugation recovery

After ultracentrifugation, the content of a tube was collected from top to bottom with 52 µL (800 µL tube) or 200 µL (5 mL tube) per fraction. We used caution to minimize disturbance to the gradient when pipetting. The recovered fractions were transferred to a 96-well plate, sealed with aluminum film, and stored at RT in the dark. To remove iodixanol and concentrate sorted liposomes, selected fractions were combined and concentrated to 50–100 µL by centrifugation (8 min at 10,000 rcf) on Amicon filtration units with 30 kD NMWL. The concentrated liposomes were diluted in Buffer X or Y to 500 µL and concentrated again for a total of 4–5 times. Optionally, sorted liposomes were treated with DNase I (Thermo Fisher Scientific) following the manufacturer's recommendation to remove DNA bricks (see **Figure S10**).

### 4. Characterization of sorted liposomes

*4a.* Agarose gel electrophoresis

Recovered fractions of a post-centrifugation gradient (5 µL each) were electrophoresed in a 3.5% agarose gel (casted with 0.05% sodium dodecyl sulfate, SDS) at 10 V/cm for 1.5 hr in a 0.5×TBE buffer containing 10 mM $MgCl_2$ and 0.05% SDS.

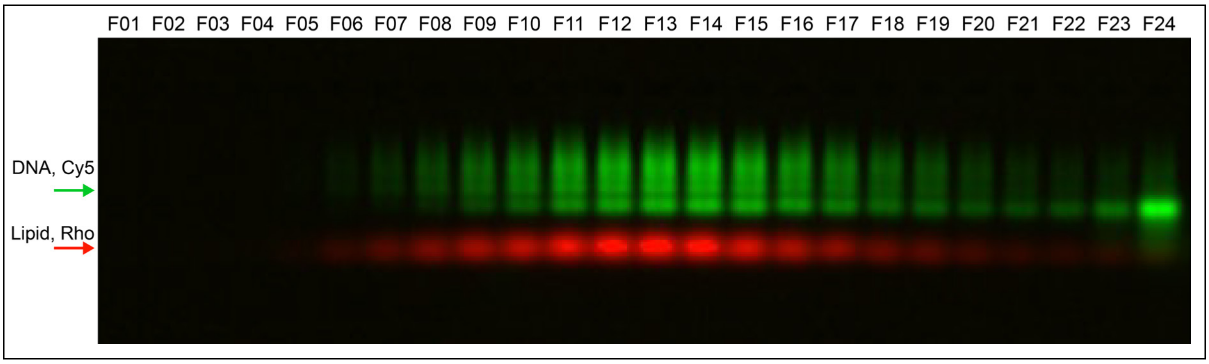

**Figure S3.** A typical SDS-agarose gel analysis showing the distribution of DNA-coated liposomes in the iodixanol gradient after isopycnic centrifugation. F01–F24 denotes the fractions collected from top to the bottom of the gradient. Pseudo-color green: Cy5-labeled DNA bricks, red: rhodamine-labeled lipids. Before running in this gel, a pool of extruded liposomes (300 pmol of total lipid, 50-nm pore size) were sorted with the help of 3PS bricks as described above. Notice that the heaviest fraction (F24) contains a large amount of DNA bricks with a negligible amount of lipids, suggesting surface saturation on most liposomes. The liposomes were lysed by SDS in the gel and running buffer, causing the lipid bands to migrate faster than the DNA-brick bands.

*4b.* Negative stain TEM study

A drop of sample (~5 µL) was deposited on a glow discharged formvar/carbon-coated copper grid (Electron Microscopy Sciences), incubated for 1–3 minutes and blotted away. The grid was then washed briefly and stained for 1 minute with 2% (w/v) uranyl formate. Images were acquired on a JEOL JEM-1400 Plus microscope (acceleration voltage: 80 kV) with a bottom-mount 4k×3k CCD camera (Advanced Microscopy Technologies). Liposome sizes were measured from electron micrographs by ImageJ (National Institutes of Health). The image analysis workflow is summarized in **Figure S4**.

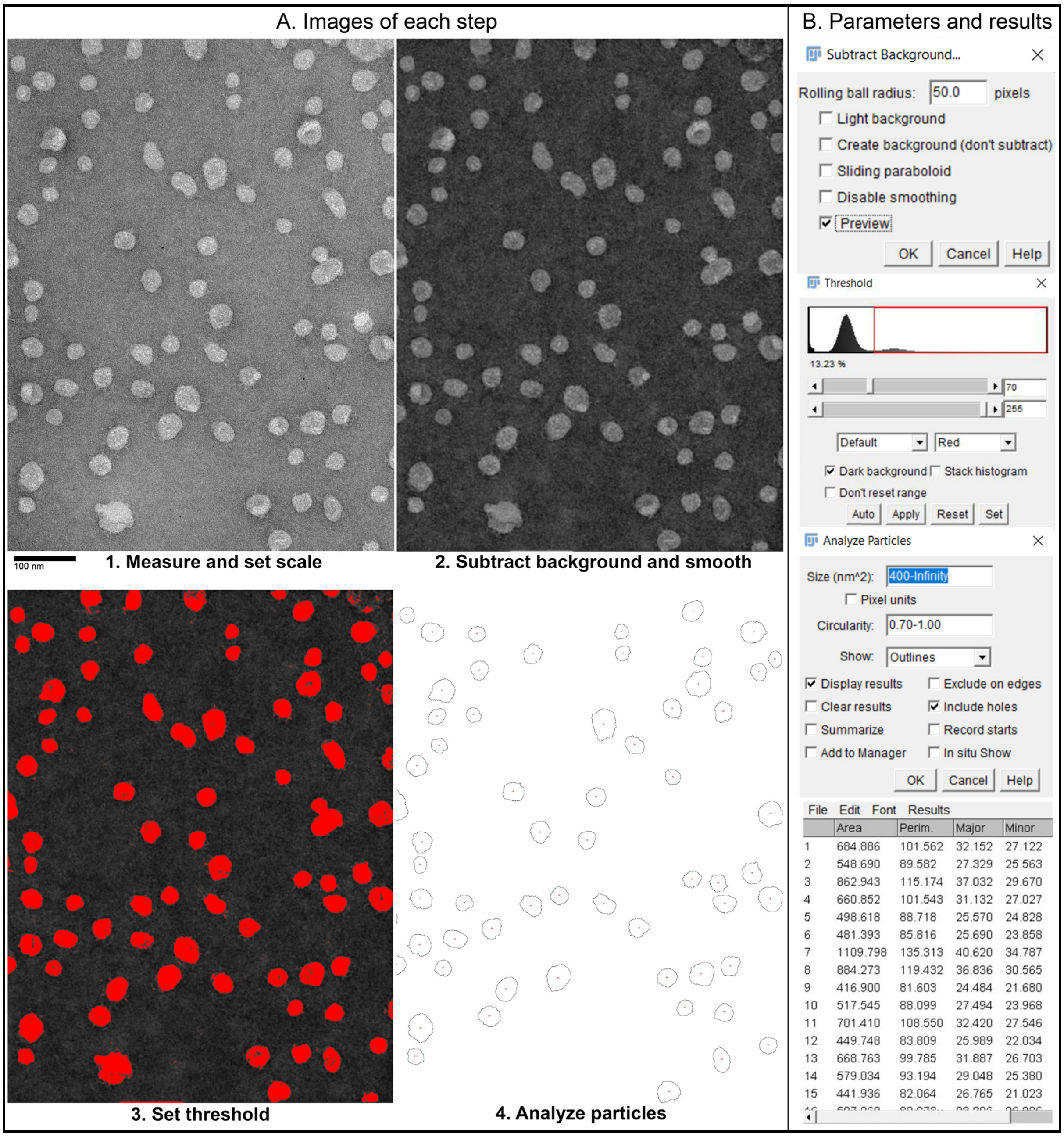

**Figure S4.** TEM image analysis pipeline to determine the diameters of sorted liposomes using the built-in functions of ImageJ. The steps are: 1. Set scale by measuring the length of the scale bar; 2. Subtract background (Rolling ball radius was set in the range of 50–150 depending on the original contrast) and smooth image (10×) for contrast enhancement and noise reduction; 3. set threshold at an appropriate value to highlight all the liposomes (holes inside are acceptable); 4. run particle analysis (circularity higher than 0.7, show outlines, include holes and display results as listed). Finally, the diameter of each liposome is calculated based on the measured area (*A*) following the equation: $\mathrm{D} = 2 \times \sqrt{\mathrm{A}/\pi}$ . Parameter setups are illustrated in panel B on the right.

*4c.* Cryo-EM imaging

A drop (3.5 μL) of liposome sample was loaded onto a glow-discharged lacey carbon film, copper, 300 mesh grids, and plunge frozen in liquid ethane using an FEI Mark III Vitrobot operating at 100% humidity, 22°C temperature, 5 s blot time and -4 force.

The grids were imaged on an FEI Talos L120C TEM equipped with a Ceta CCD camera. The images were collected at magnifications of 36K/45K/57K/92K (with the pixel size of 4.01/3.21/2.53/1.57 Å) and a dose of 50 e/Å$^2$, using a defocus range of -2 to -4 μm.

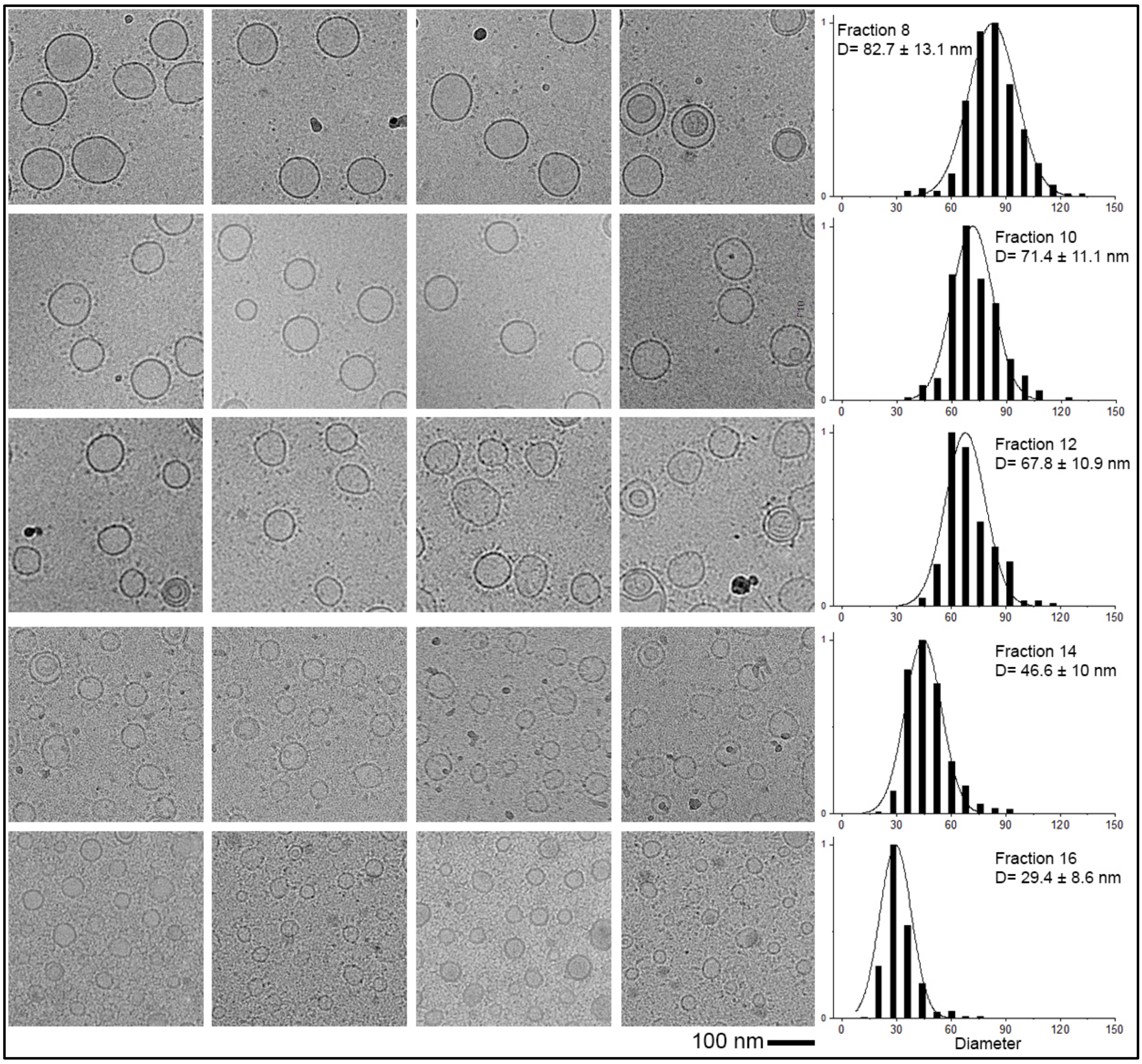

**Figure S5.** Cryo-EM images of liposomes after sorting. Representative micrographs of fractions 8, 10, 12, 14 and 16 are shown from top to bottom, together with the corresponding histograms showing the liposome size distributions. The six-helix bundle bricks are visible on the exterior surface of the liposomes. Scale bar: 100 nm.

### 5. Leakage assay (deoxyribozyme self-cleavage)

As described in Method *1a*, deoxyribozyme I-R1a (with 5'-FAM label) in Buffer X was first loaded into liposomes through a rehydration process. After sequential extrusions (Method *1b*), the liposomes were coated with 6HB bricks and sorted as described in (Method **3**). Fractions from density ultracentrifugation, as well as a control sample containing unsorted liposomes (free I-R1a pre-removal through a separate isopycnic centrifugation without DNA-brick coating), were normalized to 0.75 mM lipid concentration. A deoxyribozyme reaction buffer (DRB+) was prepared to contain 25 mM HEPES, 400 mM KCl, 6 mM $MgCl_2$ and 4 mM $ZnCl_2$, which provides the same osmotic pressure as Buffer X but with 2 mM $Zn^{2+}$ for I-R1a cleavage once mixed with the sample in a 1:1 ratio.

Three microliters of Buffer X containing 0% (for permeability test) or 14% n-octyl-β-D-glucopyranoside (OG, for liposome lysis) were added to 9 µL of each sample (fraction 6, 8, 10, 12, 14, 16, 18, and unsorted), then mixed with 12 µL DRB+ and incubated at 37°C for 12 hr. After incubation, samples were mixed with 16 µL denaturing loading buffer (90% formamide, 10 mM NaOH, 1 mM EDTA, 0.1% Xylene Cyanole FF) and boiled for 3 min. Samples (10 µL each) were electrophoresed in a 12% urea polyacrylamide gel containing 0.1% SDS in 1×TBE buffer with 0.1% SDS at 10 V/cm for 1.5 hours (**Figure 2**).

### 6. ATG7/ATG3 catalyzed GL1 lipidation assay

*6a.* Lipidation reaction

Protein expression and membrane-curvature dependent lipidation reactions were performed as described previously[1-3]. Purified ATG7 (1.5 µM), ATG3 (2.5 µM) and human GABARAP L1 (GL1, 8 µM) were mixed with unsorted or sorted liposomes (composition B, 1 mM total lipid) in the presence of 1 mM dithiothreitol (DTT) and 1 mM ATP in SNH buffer (50 mM Tris at pH 8, 100 mM NaCl and 1 mM $MgCl_2$) and incubated at 30 °C for 90 min. The reaction was stopped by 4× SDS-PAGE loading buffer and boiled at 90°C for 5 min. Electrophoresis was performed in precast 10% Bis-Tris gels (Novex, Thermo Fisher Scientific) running in 1×MES SDS Running Buffer (NuPAGE, Thermo Fisher Scientific) at 180 V (18V/cm) for 60 min. The proteins were visualized with Coomassie blue stain following the manufacturer's instruction (Imperial Protein Stain, Thermo Fisher Scientific).

*6b.* Immunoblotting

After electrophoresis, samples were transferred onto a PVDF membrane (Amersham, GE Healthcare), blocked with 5% BSA and probed with anti-GL1 (1:1000, Cell Signaling Technology clone D5R9Y) antibody (Sigma) in 2.5% BSA. HRP-conjugated anti-mouse (NA931) and anti-rabbit (NA934) secondary antibodies were purchased from Amersham, GE Healthcare. See **Figure 3** and **S12** for results.

### 7. SNARE-mediated liposome fusion assay

*7a.* Plasmid constructs and protein purification

The vectors encoding full-length t-SNARE complex including rat Stx1A and mouse 6×His-SNAP25 (plasmid pTW34) and 6×His-SUMO-VAMP2 (plasmid pET-SUMO-VAMP2), were transfected into the BL21-Gold (DE3) *E. coli* strain (Agilent Technologies; Cat# 230132) and purified as previously described[4]. Briefly, bacteria carrying SNARE plasmids were cultured in 2 L LB media at 37 °C until $OD_{600}$ reached 0.7, induced by 1 mM isopropyl β-D-thiogalactoside, and cultured for additional 3 hr at 37 °C. The pelleted cells were resuspended in breaking buffer (25 mM HEPES pH 7.4, 400 mM KCl, 10% glycerol, 4% Triton X-100, 1 mM TCEP, protease cocktail inhibitors) and lysed by cell disruptor (Avestin) with 3–5 passages at ~15,000 psi. The cell lysate was clarified by centrifugation at 40,000 rpm for 30 min; the supernatant was collected and incubated with nickel-NTA agarose (Qiagen) for 4 hr to overnight at 4°C. t-SNARE bound beads were rinsed with 25 mM HEPES pH 7.4, 400 mM KCl, 10% glycerol, 1% (w/v) OG, 1 mM TCEP. t-SNARE proteins were eluted off the beads by elution buffer (25 mM HEPES pH 7.4, 400 mM KCl, 10% glycerol, 1% OG, 1 mM TCEP, 400 mM imidazole). 6×His-SUMO tags on VAMP2 were cleaved by SUMO protease.

*7b.* Proteoliposome preparation

SNARE proteins were reconstituted into liposomes at physiologically relevant densities, with protein:lipid ratio at 1:200 or 1:400 for v-SNARE liposome and 1:400 for t-SNARE liposomes. A vacuum-dried lipid film was dissolved in the reconstitution buffer (25 mM HEPES pH 7.4, 140 mM KCl, 0.2 mM TCEP, 10% glycerol, 1% OG) and mixed with SNARE proteins. OG-free reconstitution buffer was added to reach a final OG concentration of 0.33%. Detergent was then removed in a Slide-A-Lyzer dialysis cassette (Thermo Fisher Scientific) against 4 L of

OG-free reconstitution buffer at 4°C overnight. Proteoliposomes were separated in a Nycodenz (Progen Biotechnik) density gradient via centrifugation[5]. For t-SNARE liposomes, centrifugation was done in an SW60-Ti rotor (Beckman Coulter) at 55,000 rpm for 3hr 40min at 4 °C; for v-SNARE liposomes, centrifugation was done in an SW55-Ti rotor (Beckman Coulter) at 48,000 rpm for 4 hr at 4 °C. The enriched proteoliposomes were collected from 0/30% Nycodenz interface. These proteoliposomes were sorted as described in Methods **3** and analyzed by negative-stain TEM (Method *4b*) and SDS-PAGE (**Figure S14**). The v-SNARE concentrations of proteoliposomes were determined using VAMP2 concentration standards by densitometry (ImageJ). Lipid concentrations of v-SNARE liposomes were determined by rhodamine absorbance at 574 nm.

*7c.* Lipid mixing assay

A typical fusion reaction happened between 5 µL of v-SNARE liposomes labeled with a pair of FRET dyes (donor: NBD-DOPE, acceptor: Rhodamine-DOPE) and 45 µL of unlabeled t-SNARE liposomes[5] (**Table S2**), with a total lipid concentration of 3 mM. These 50 µL mixtures were pre-incubated at 4 °C for 2 hr for trans-SNARE complex assembly, before being transferred to a Falcon 96-well plate with a black skirt and clear flat bottom and heated to 37 °C. NBD fluorescence was monitored at emission/excitation of ~535/460 nm every 1 min for 2 hr by Synergy H1 Hybrid Multi-Mode Reader (BioTek Instruments). At the end of the 2-hr reaction, 10 µL of 20% Triton X-100 was added and fluorescence was recorded for another 30 min to obtain the maximum fluorescence.

**Results.**

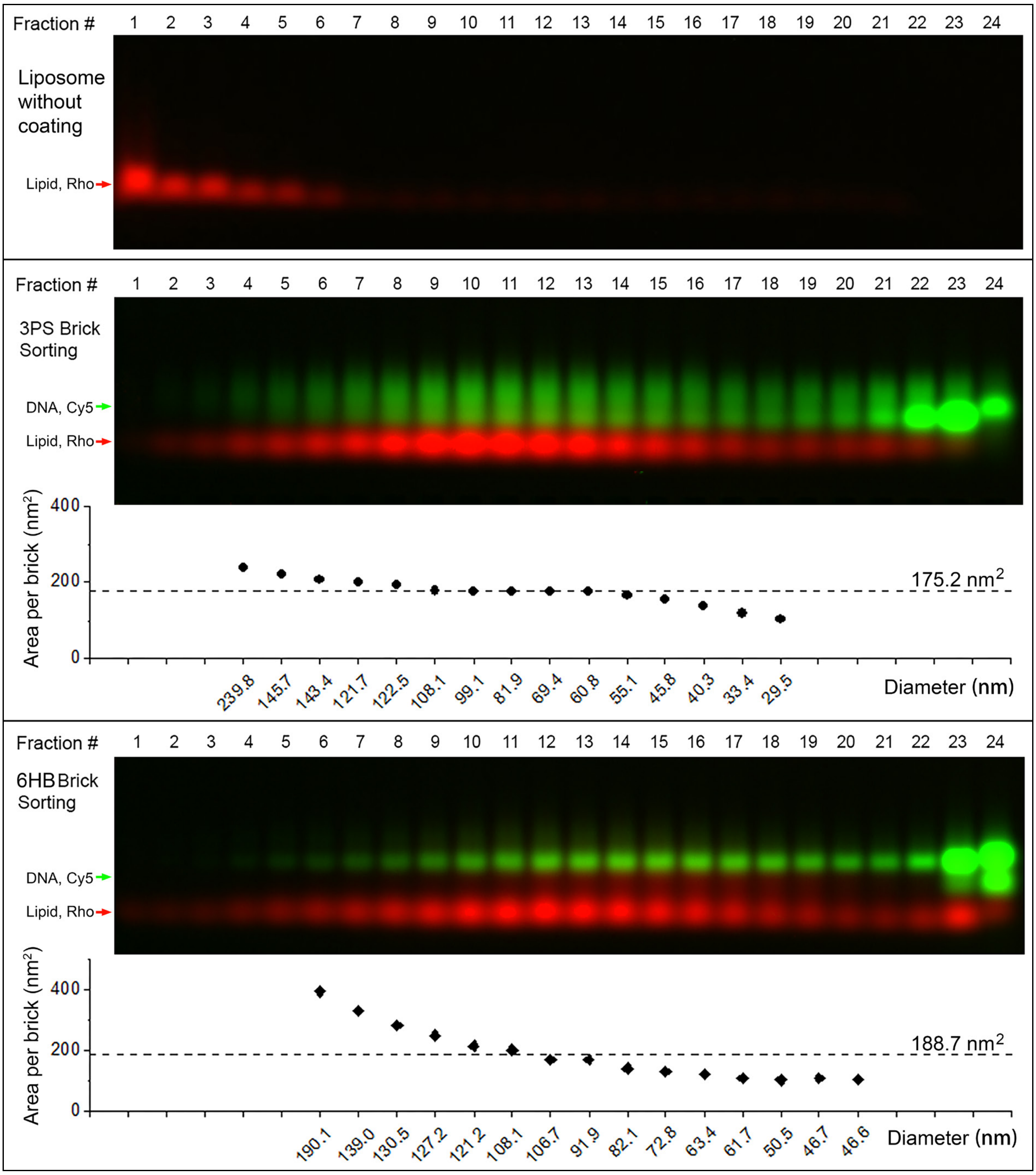

**Figure S6.** Uncoated (top) and 3PS-(middle)/6HB-(bottom) brick coated liposomes after ultracentrifugation in iodixanol density gradients (Method **3**). Fractions are electrophoresed in the same SDS-agarose gel (Method *4a*). Liposomes to be sorted consist of a 1:1 (molar ratio of total lipid) mixture of extruded (through 50-nm pores) and sonicated liposomes. The average surface area occupied by each brick is calculated based on lipid:DNA ratio estimated from the band intensities. On average, each brick occupied ~200 $nm^2$ of membrane surface. Bricks bound stronger to smaller liposomes, presumably because of more lipid packing defects in the highly curved membranes.

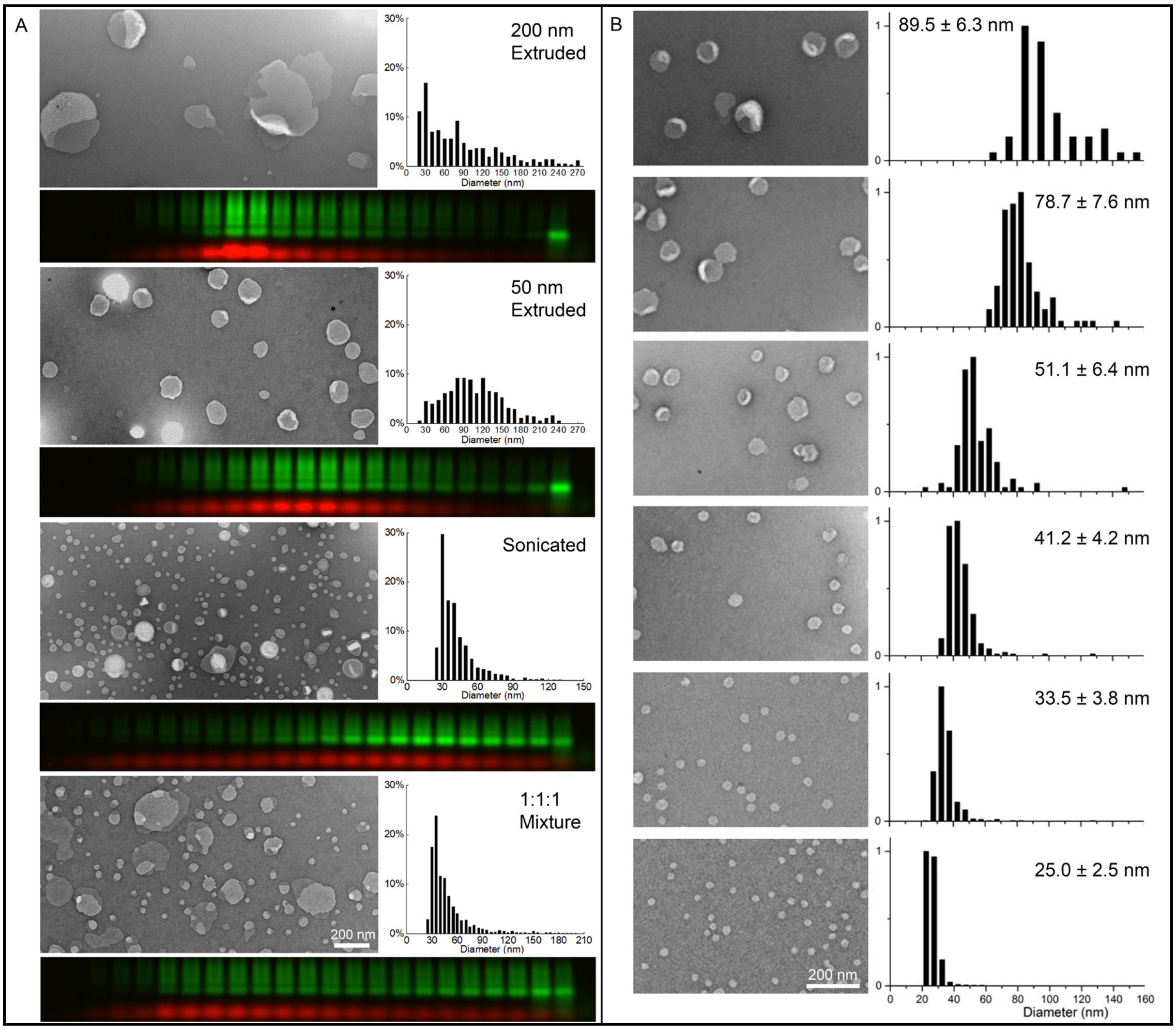

**Figure S7.** DNA-brick assisted sorting of liposomes of different origins and size distributions. (A) Various heterogeneous liposomes (extruded liposomes, sonicated liposomes, and their mixture) and the sorting results (analyzed by SDS-agarose gel electrophoresis). For each sample, a representative TEM image, a histogram of liposome diameters, and a pseudo-colored agarose gel (red: rhodamine-labeled lipid, green: Cy5-labeled DNA) containing 3PS-brick assisted sorting products are shown. The distributions of rhodamine fluorescence within the gradients reflect the spectra of liposome size before sorting. The diameter histogram of the 1:1:1 (molar ratio of total lipid) liposome mixture is similar to that of the sonicated liposomes because of a dominant population of <40-nm liposomes in the 1:1:1 mixture. (B) Representative TEM images and diameter histograms of liposomes sorted from the 1:1:1 mixture with the help of 3PS bricks. Scale bar: 200 nm.

Note: Reconstituted liposomes can be sorted successfully as well (**Figure S14**).

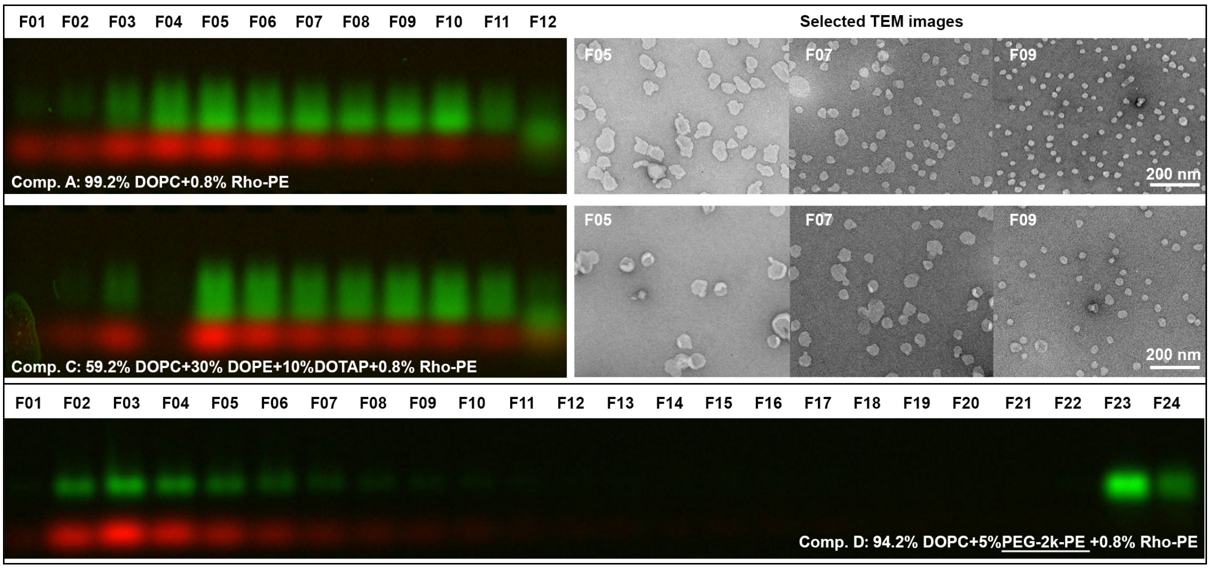

**Figure S8.** DNA-brick assisted sorting of liposome of different lipid compositions. Top row: small scale (1×, **Table S3**) sorting of liposomes of Composition A (**Table S2**). A pseudo-colored SDS-agarose gel (red: rhodamine-labeled lipid, green: Cy5-labeled DNA) containing density-gradient fractions is shown to the left of the representative TEM images (scale bar: 200 nm). Middle row: same as top row, but with Composition C. The changes in lipid charge do not have a major impact on sorting results. Bottom row: Attempted large scale (10×, **Table S3**) sorting of liposomes with Composition D. The PEGylated lipids hamper DNA-brick coating and thus negatively impact sorting.

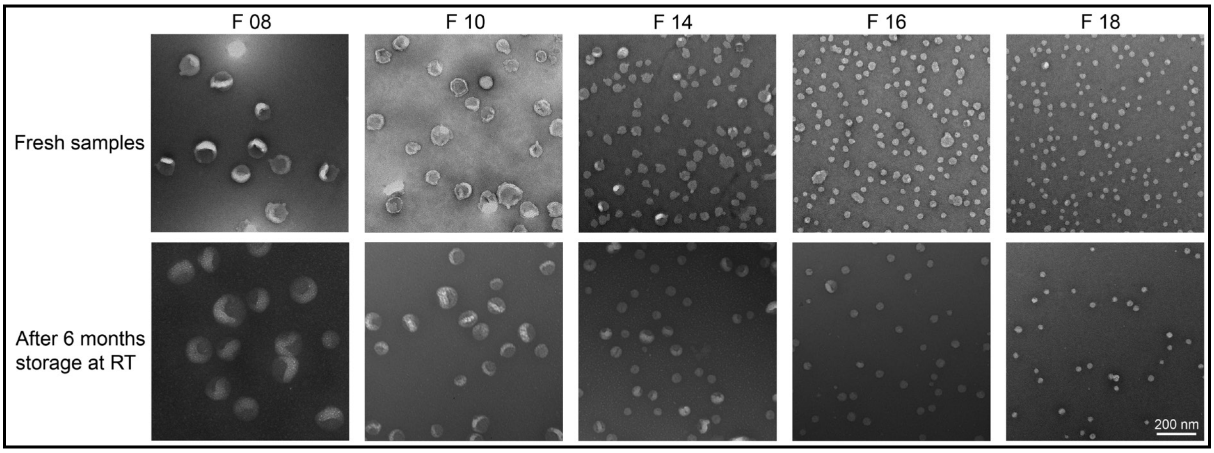

**Figure S9.** Stability of the sorted liposomes. Top row: TEM images of freshly sorted liposomes with 3PS-brick coating. Bottom row: TEM images of the same fractions after 6-month storage at room temperature. Scale bar: 200 nm.

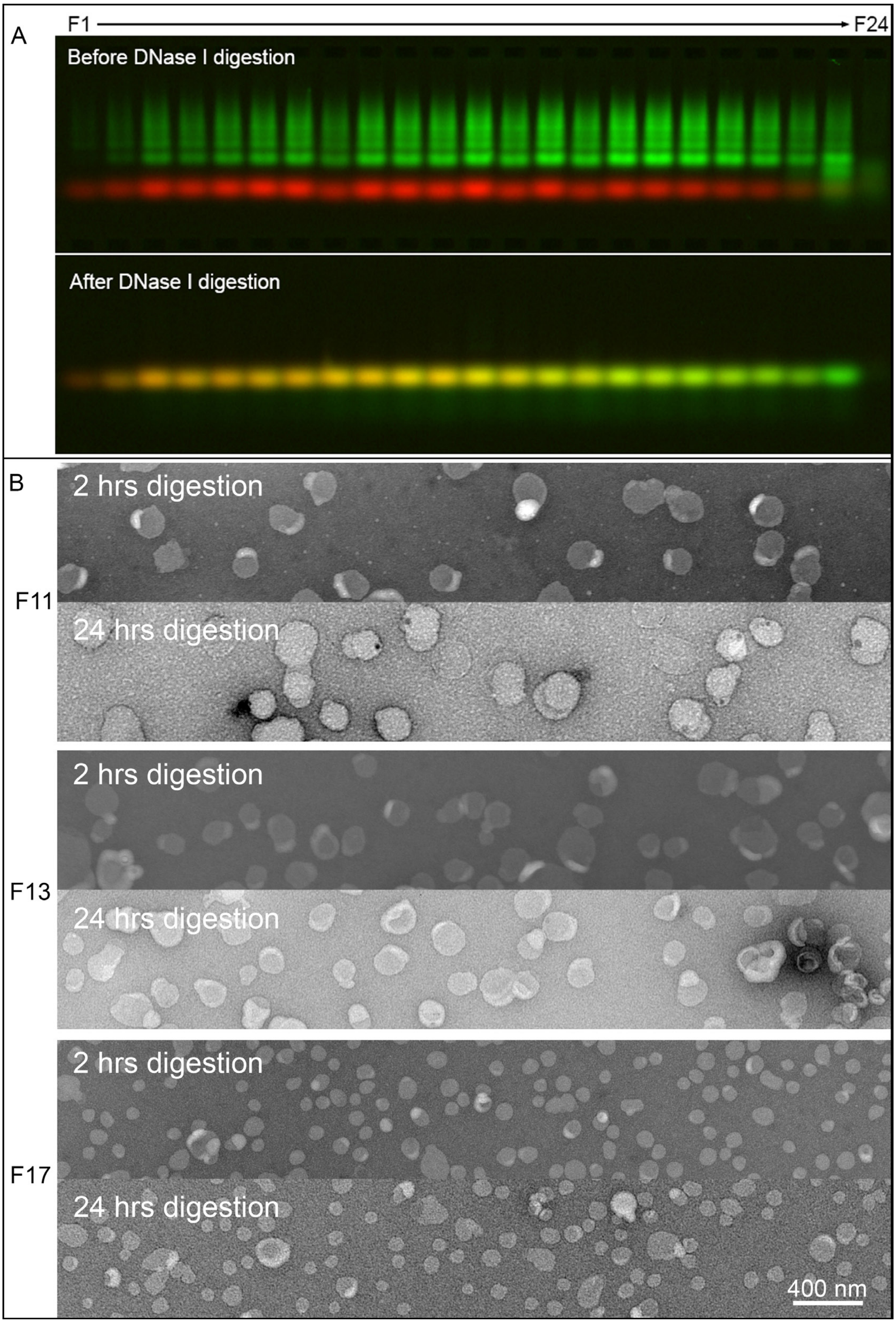

**Figure S10.** Enzymatic removal of DNA bricks from sorted liposomes. (A) SDS-agarose gel analyses of sorted liposomes before (top) and after (bottom) nuclease treatment. One unit of DNase I is added to 100 µL of fractionated liposomes (coated by 3PS brick) and incubated at 37°C for 24 hours. Pseudo-color green: Cy5-labeled DNA, red: rhodamine-labeled lipid. (B) TEM images of selected fractions (F11, F13 and F17) showing liposomes treated by DNase I for 2 or 24 hours.

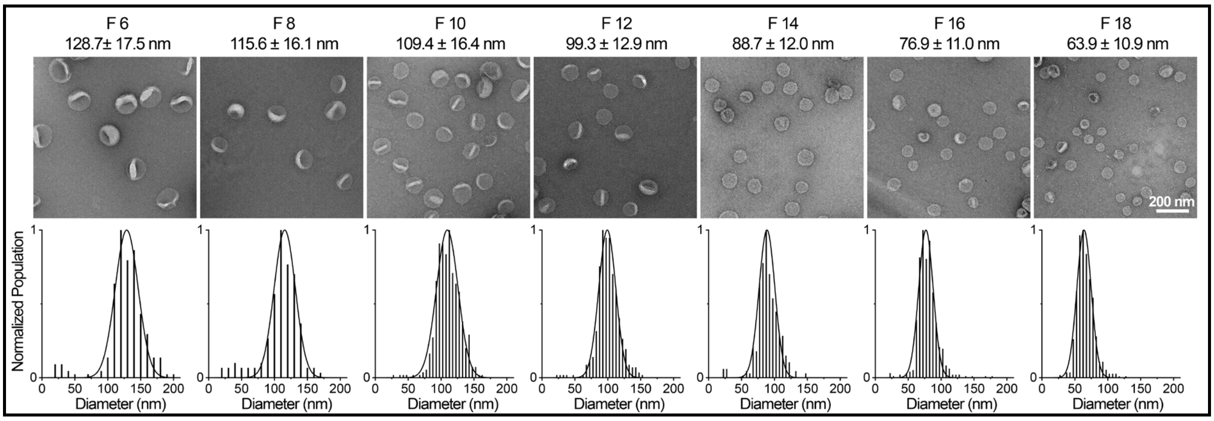

**Figure S11.** 6HB-brick assisted sorting of deoxyribozyme encapsulating liposomes. For sorted liposomes in each fraction, a representative TEM image is shown on top of the corresponding histogram of diameters. Liposomes contain deoxyribozyme I-R1a and are sorted with the help of 6HB bricks. Scale bar: 200 nm.

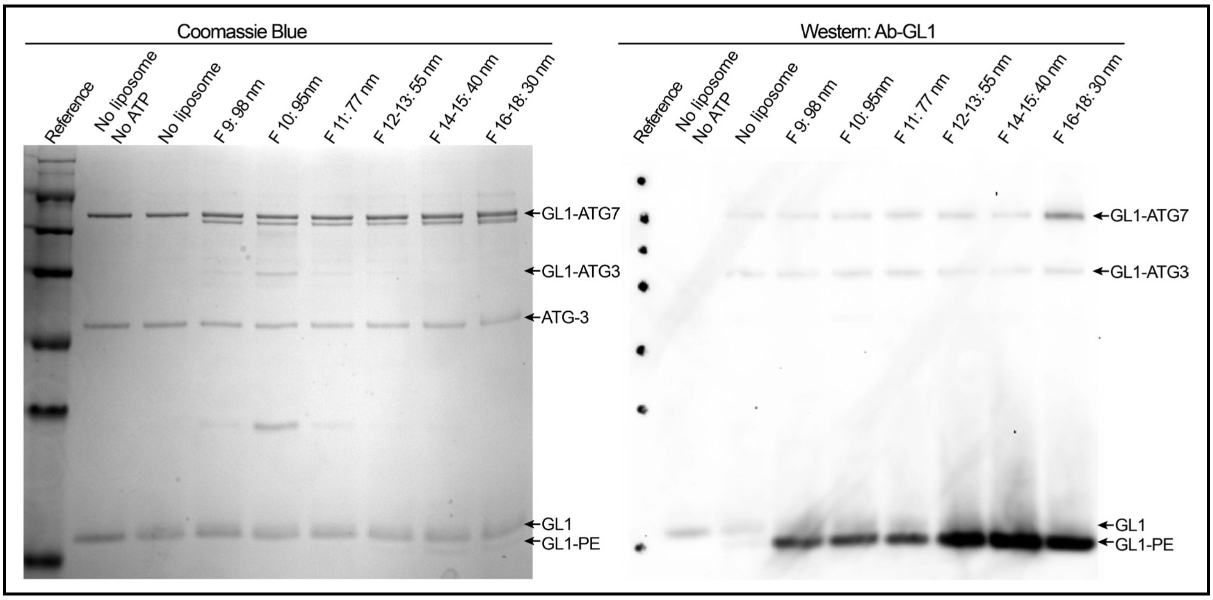

**Figure S12.** Curvature dependency of ATG7/ATG3 catalyzed GL1 lipidation. Typical SDS-PAGE and western blot analyses of GL1 lipidation reactions are shown on the left and right panels, respectively. Reaction conditions are summarized in Method **6**.

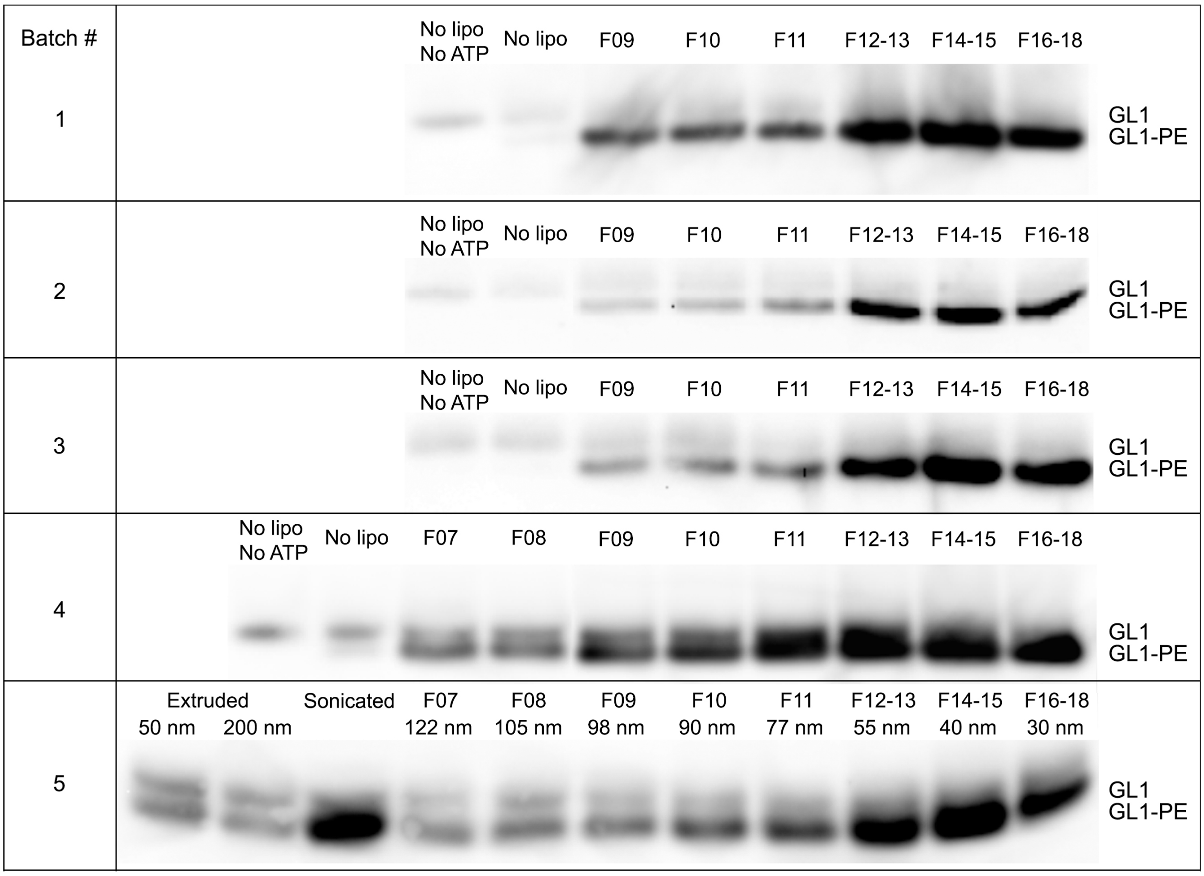

**Figure S13.** ATG7/ATG3 catalyzed GL1 lipidation using sorted liposomes from five separate preparations (batch 1–5) analyzed by western blot.

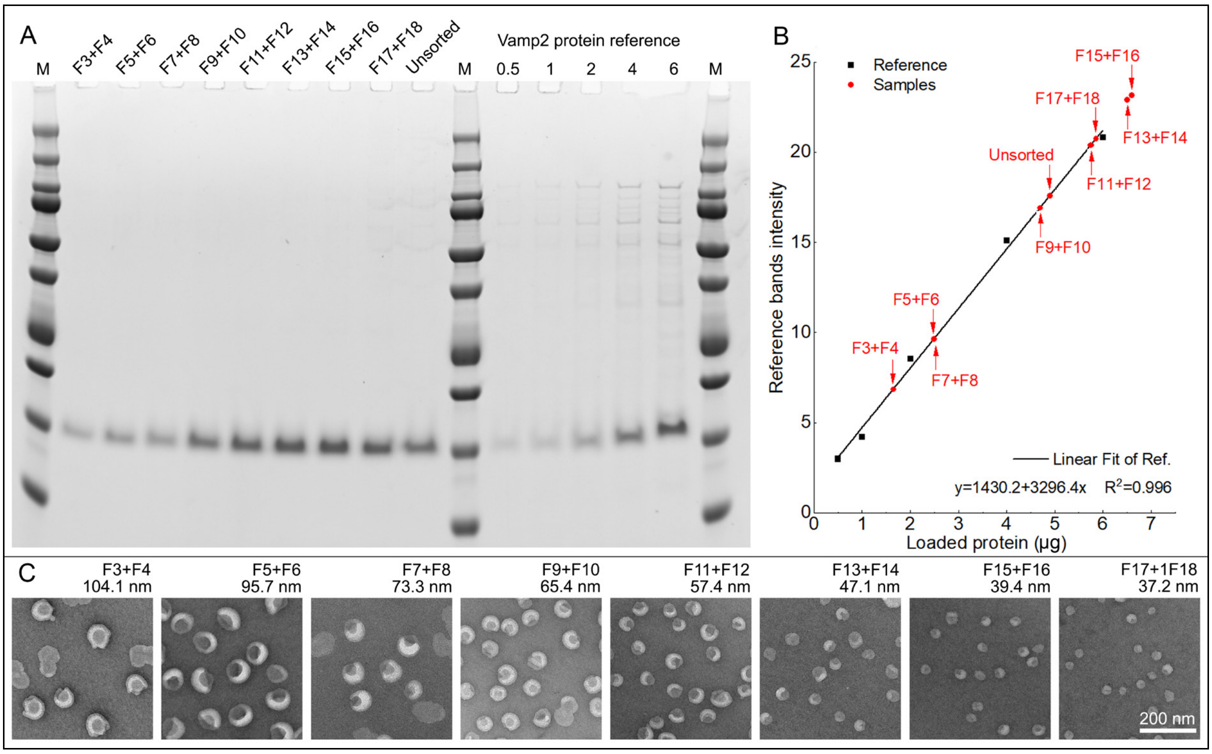

**Figure S14.** Characterizations of proteoliposomes containing VAMP2. (A) and (B) Quantification of VAMP2 protein in reconstituted proteoliposomes before and after sorting. Liposomes reconstituted with VAMP2 are analyzed by SDS-PAGE alongside with protein concentration references (A). A linear regression of reference band intensity on the mass of proteins generates a calibration curve (B), which is used to calculate the amount of VAMP2 in the proteoliposomes before and after sorting. (C) Representative TEM images of VAMP2-containing liposomes after sorting. Fraction numbers and mean diameters on top of the corresponding TEM images. Scale bar: 200 nm.

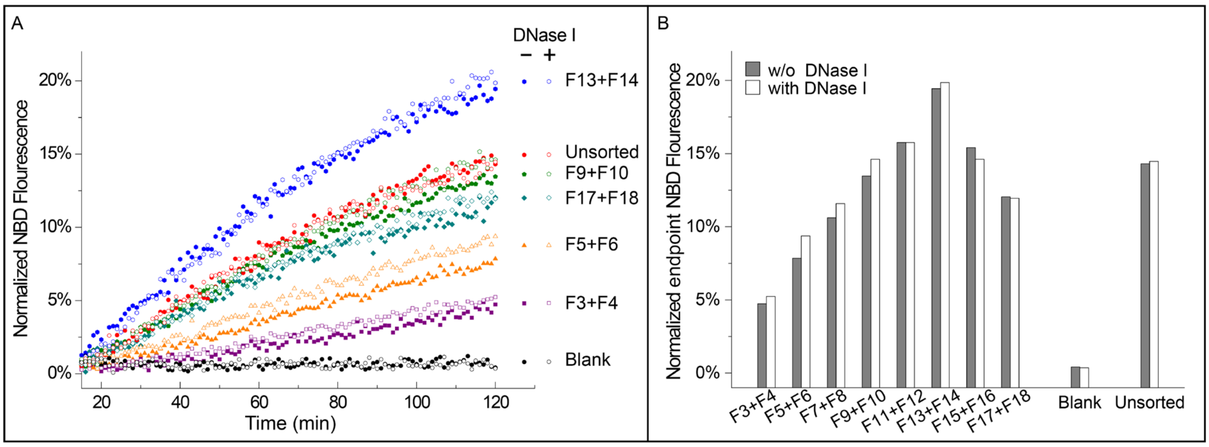

**Figure S15.** Effect of membrane-bound DNA bricks on fusion assay. (A) NBD fluorescence traces showing the lipid mixing kinetics between unsorted t-SNARE liposomes and unsorted or sorted v-SNARE liposomes with or without DNase I digestion (1 U/10 μL, 37°C, 2 hours). (B) NBD fluorescence after 2 hours of fusion reactions (Method **7**, fluorescence traces shown in (A)). See **Figure 4** or **S14** for the mean diameters of sorted liposomes in each fraction (Fx).

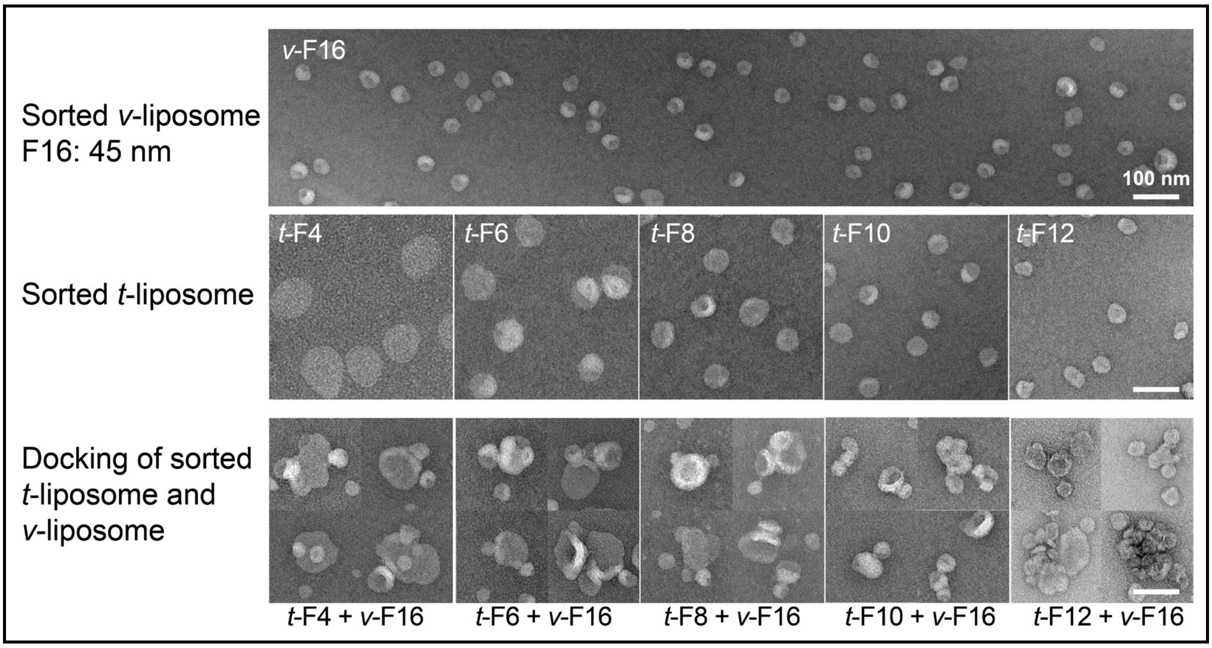

**Figure S16.** Liposome docking in the pre-incubation period visualized by negative-stain TEM. Top row: a homogeneous population of v-SNARE liposomes after sorting (fraction 16, mean diameter: 45 nm). Middle row: t-SNARE liposomes sorted into five homogeneous populations (fractions 4, 6, 8, 10 and 12). Bottom row: incubating v-SNARE liposomes (45-nm mean diameter) and t-SNARE liposomes of various homogeneous sizes for 2 hours at 4°C (i.e., pre-incubation, see Method **7**) results in vesicle clusters, suggesting docking between the two proteoliposome species. Scale bars: 100 nm.

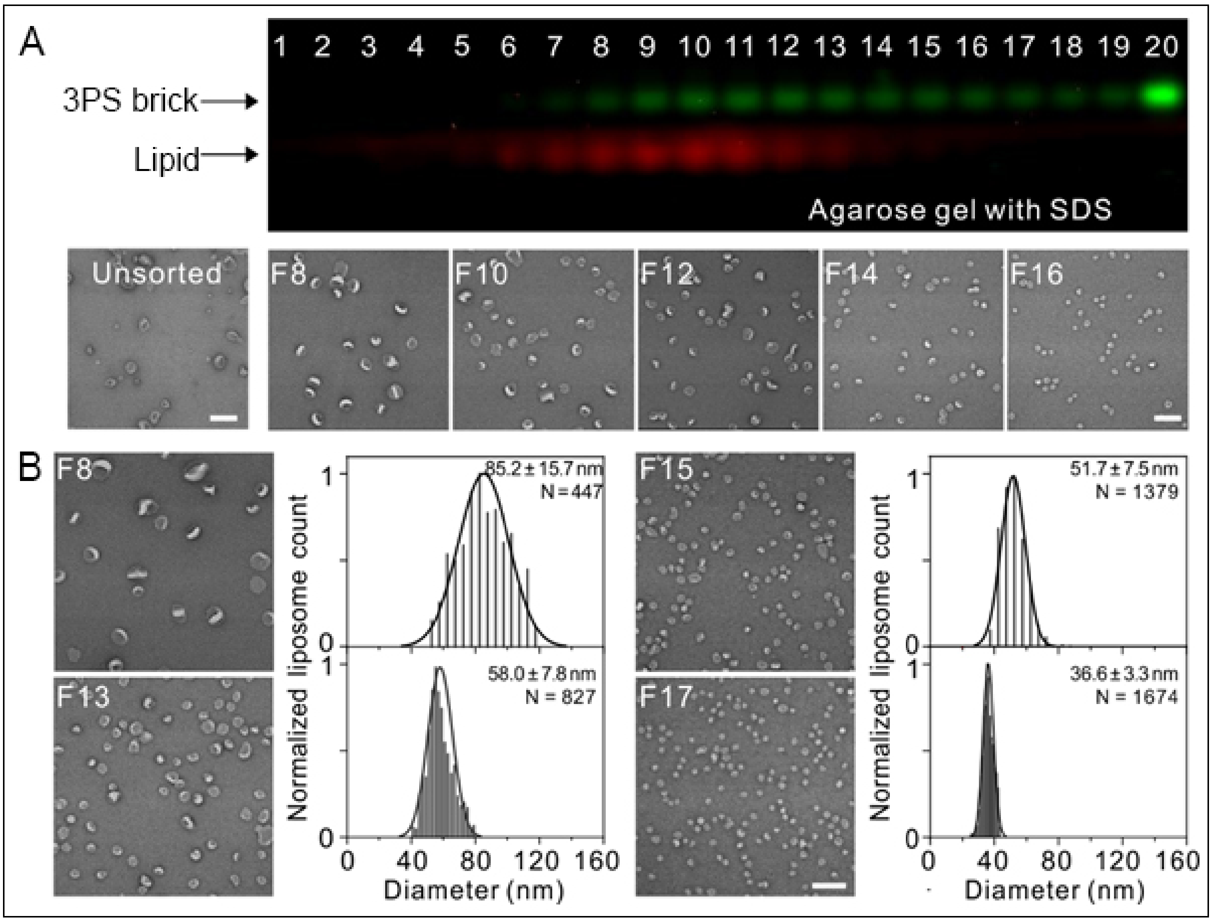

**Figure S17.** Sorting experiment reproduced at Fudan University. (A) 3PS-brick-assisted sorting of liposomes (extruded through 50-nm filters, 0.4 µmol of total lipids) analyzed by SDS-agarose gel electrophoresis (top) and negative stain TEM (bottom). Fractions are numbered sequentially from F1 to F20 from top to bottom of the gradient. (B) The size distribution of sorted liposomes in selected fractions (F8, F13, F15 and F17) measured from negative-stain TEM images. Histograms are fitted to Gaussian curves. Fitted means and standard deviations (mean ± SD) of liposome diameters and sample sizes (N) are noted with the corresponding histograms. Scale bar: 200 nm.
